# RIPK4 driven by TP53 mutations promotes resistance to redox stress of CRC by phosphorylating MTHFD1

**DOI:** 10.1101/2024.08.06.606759

**Authors:** Long Yu, Sha Zhou, Yan-Bo Xu, Zhong-Jin Zhang, Xiao-Man Cheng, Chi Zhou, Wei-Hao Li, Jia-Hua He, Qing-Jian Ou, Jia-Yi Qin, Yu-Jing Fang, Jian-Hong Peng, Jun-Zhong Lin, Bo Lin, Zhen-Lin Hou, Zhi-Zhong Pan

## Abstract

This study investigates advanced colorectal cancer (CRC), focusing on its tendency for distant metastasis and chemotherapy resistance. It highlights the importance of PANoptosis, a cell death pathway, and the role of the Receptor-interacting serine/threonine-protein kinase (RIPK) family in tumor progression. RIPK4’s tissue-specific functions in cancer cell behavior are emphasized, including its influence on invasion, migration, and oxidative stress resistance. The study reveals the critical balance of reactive oxygen species (ROS) in cancer cells, linked to antioxidant defenses and NADPH production for survival. A key finding is the connection between TP53 mutations in CRC and increased RIPK4 expression, which enhances MTHFD1 phosphorylation, boosts NADPH production, reduces ROS, and promotes resistance to PANoptosis, leading to metastasis. The research identifies the molecular basis of CRC metastasis, showing how RIPK4 regulates MTHFD1 to resist PANoptosis, offering new therapeutic targets for metastatic CRC and potential improvements in patient outcomes.

## Introduction

Eighty-three percent of colorectal cancer (CRC) patients present with advanced stages, with approximately 44 percent having distant metastases at diagnosis(*1*). This advanced state significantly challenges clinical treatment, as evidenced by a five-year survival rate below 20 percent for these patients, even after comprehensive treatment(*2, 3*). The high mortality rate of metastatic CRC is primarily attributed to its poor prognosis(*4, 5*). A key factor contributing to this is the resistance and perturbation of PANoptosis, a programmed inflammatory cell death pathway that integrates features of pyroptosis, apoptosis, and necroptosis (*6–8*). During metastasis, tumor cells must overcome various cell death mechanisms, including PANoptosis(*6, 9*). However, the mechanisms by which tumor cells regulate PANoptosis and metastasis remain largely unexplored.

The Receptor-interacting serine/threonine-protein kinase (RIPK) family comprises seven kinases that share a homologous kinase domain at the N-terminus. RIPKs are integral to immune responses, inflammation, and cell death, with each member playing specific roles based on their functional domains(*10*). Structurally, RIPK1, -2, -4, and -5 have intermediate domains. However, only RIPK1 contains the RIP homotypic interaction motif (RHIM) in this domain, allowing interaction with RIPK3’s RHIM at the carboxyl terminus. RIPK4 and RIPK5 contain ankyrin repeats at their C-terminus(*11*). While RIPK1-3 are well-studied, less is known about RIPK4-6. In the existing research, RIPK4 is considered to be the binding partner of PKC β and PKC δ proteins, and it is mainly involved in the key regulatory factors of keratinocyte differentiation, skin inflammation and skin wound repair(*12, 13*). In pancreatic cancer, overexpression of RIPK4 promotes invasion and migration of tumor cells by activating the RAF1/MEK/ERK pathway(*14*). RIPK4 can also upregulate VEGF-A through NF-kB pathway to promote the invasiveness of bladder urothelial carcinoma cells(*15*). But, the expression of RIPK4 is tissue specific, and its biological function may be different(*16–19*).

Tumor cells undergoing metastasis often exhibit increased levels of reactive oxygen species (ROS) levels compared with primary lesions(*20–22*). Optimal ROS levels are crucial for signaling and cellular functions in cancer cells(*23, 24*). Excessive ROS levels can induce cytotoxicity, DNA damage, and cell death(*25*). To prevent excessive oxidative stress and maintain favorable redox homeostasis, tumor cells have evolved a complex antioxidant defense system that strategically regulates multiple antioxidant enzymes such as catalase, glutathione reductase, and antioxidant molecules. Antioxidant molecules depend on the production of nicotinamide adenine dinucleotide phosphate (NADPH), which is used to maintain reduced glutathione (GSH) and thioredoxin (TRX)(*26–28*). NADPH homeostasis is mainly regulated by several metabolic pathways and enzymes, including NAD kinase (NADK), the pentose phosphate pathway (PPP), folate mediated one-carbon metabolism, malic enzyme (ME), NADP dependent isocitrate dehydrogenase (IDH1 and IDH2), glutamine metabolism and fatty acid oxidation (FAO)(*20, 29, 30*). One-carbon metabolism is a universal metabolic process in cells in which organic groups containing one-carbon atoms are transferred to participate in nucleotide biosynthesis, providing methyl groups for all biological methylation reactions. This process is particularly important for cancer metabolism. In addition, the reducing substance NADPH is produced(*31, 32*). Targeting one-carbon metabolism for cancer treatment has been used for cancer treatment since 1948. Early anti-folate chemotherapies such as methotrexate remain effective cancer treatments today. However, side effects are common and may be severe. Methylenetetrahydrofolate Dehydrogenase, Cyclohydrolase And Formyltetrahydrofolate Synthetase (MTHFDs) are the key enzymes involved one-carbon metabolism. The importance of MTHFDs in normal growth and development make the upregulation of these enzymes valuable to cancer cells as well. MTHFD1, a member of the MTHFD family, has been found to be an important predictor of prognosis in several cancers, and might be a promising therapeutic target in cancer.

Approximately 60% of CRCs cases exhibit a wide range of TP53 mutations, which are associated with increased metastasis and poor prognosis (*33, 34*). Recent evidence suggests that some TP53 mutations may endow mutant p53 with oncogenic properties, actively driving cancer progression(*35–40*). Our study revealed that RIPK4 expression, driven by TP53 mutation, was upregulated in CRC. This elevation in RIPK4 promotes the phosphorylation of MTHFD1, enhances NADPH production, reduces ROS levels, fosters resistance to PANoptosis and promotes distant metastasis. These findings not only elucidate a novel mechanism of metastasis driven by TP53 mutation but also underscore the potential of targeting RIPK4 in the treatment of metastatic CRC.

## Results

### Tumor cell death during suspension

First, we aimed to understand the changes in tumor cells that are detached from the extracellular matrix during suspension. We performed RNA-seq on CRC cell SW620 after 72 hours of suspension. As shown in **Supplementary Fig. S1A**, suspended CRC cells exhibit biological changes due to hypoxia and changes in extracellular matrix components. This suggests that tumor cells may experience oxidative stress and cell death during suspension. Next, we used two cell death dyes, YP1 and PI, to observe the type of death experienced by the suspended tumor cells. YP1 positive cells indicated apoptosis or necroptosis, and PI positive cells indicate necroptosis, pyroptosis, or ferroptosis(*7*). Both SW620 and LoVo cells showed an increase in YP1 and PI signals during suspension, particularly the PI signal. We also observed classical balloon-like cell death structures in suspended LoVo cells (**Supplementary Fig. S1B and Supplementary Fig. S1C**). These results suggested that the suspended tumor cells not only experienced apoptosis, but also other cell deaths. In the vicinity of 70-100 kDa, we observed significant changes in protein content and identified the relevant proteins using mass spectrometry (**Supplementary Fig. S1D**). Among the identified proteins, we were more interested in the RIPK4 protein kinase, which was verified by western blotting (**Supplementary Fig. S1D**). RIPK4 protein increased significantly in the suspended tumor cells, and its protein mass spectrum is shown in **Supplementary Fig. S1E**.

### High expression of RIPK4 is associated with poor prognosis

We explored the expression of RIPK4 in CRC using several Gene Expression Omnibus (GEO) datasets. In the GSE41258 dataset containing polyps, primary tumors and liver metastases, the expression of RIPK4 gradually increased compared to that in normal tissue. Elevated RIPK4 expression was also observed in primary tumors in the GSE9348 and GSE20916 datasets. In the GSE28702 dataset, the liver metastasis of CRC showed higher expression levels (**Fig. 1A**). Therefore, we suspected that RIPK4 plays a role in the progression and metastasis of CRC. Next, we used IHC to verify the expression of RIPK4 in normal tissues, orthotopic tumors and liver metastases. As shown in **Fig. 1B**, RIPK4 protein expression gradually increased in the tissues of two paired patients with liver metastasis, and the IHC scores are shown in **Fig. 1C**. We also observed the relationship between RIPK4 expression and prognosis in the TCGA-COADREAD (The Cancer Genome Atlas Program) dataset. As shown in **Fig. 1D**, patients with high RIPK4 expression had poor overall survival, disease-specific survival, disease-free survival, and progression-free survival. Consistent results among the 56 CRC samples collected by SYSUCC (**Fig. 1E**). For the analysis of tissue protein content, we collected 12 pairs of CRC tumor tissues and paired normal tissues and validated the RIPK4 protein content by western blotting. As shown in the **Fig. 1F**, RIPK4 protein was highly expressed in the tumor tissues of 9 of the 12 patients. The expression of RIPK4 protein in two patients with liver metastasis CRC was consistent with the results from the public database (**Fig. 1G**).

**Fig. 1.**
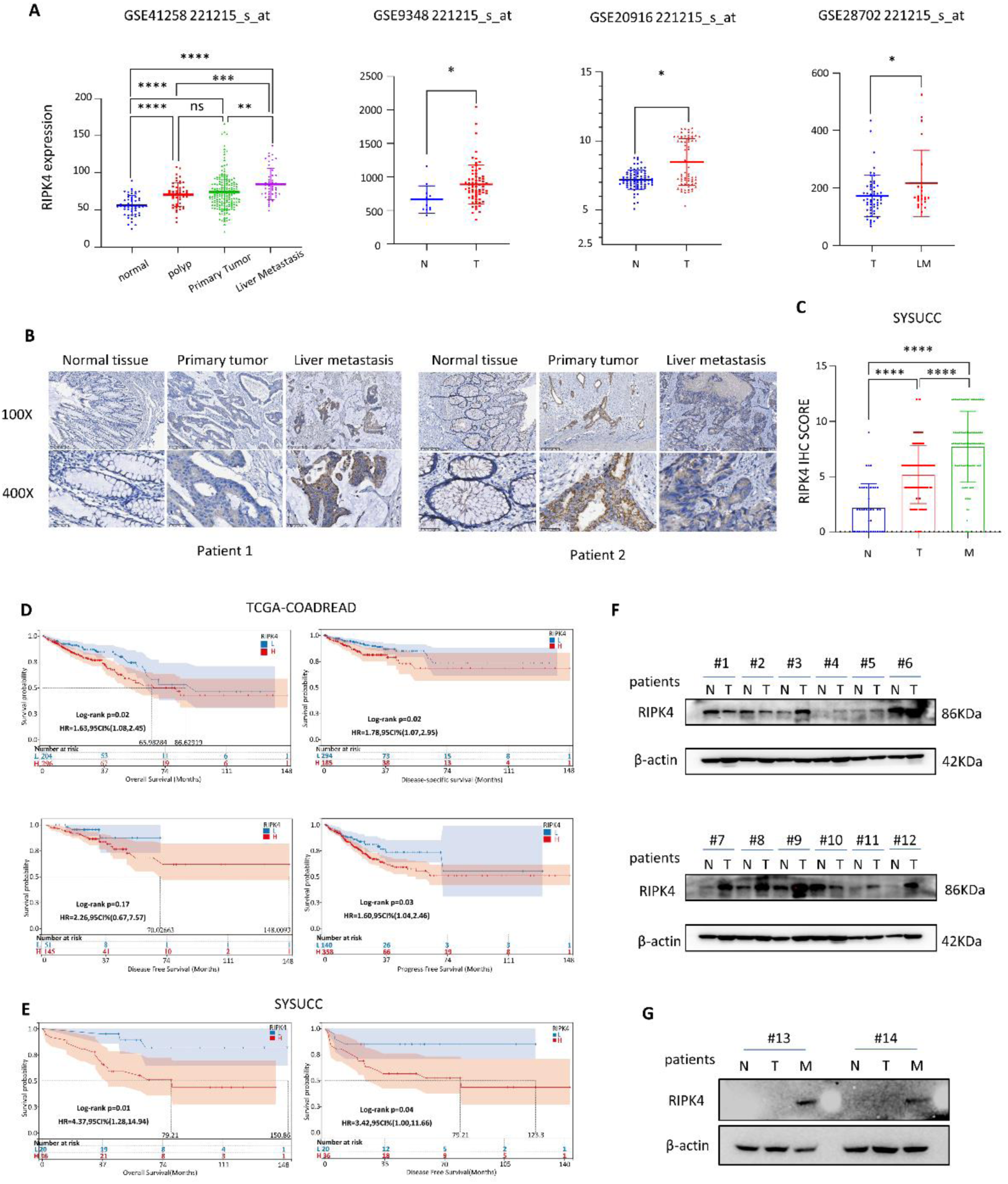
High expression of RIPK4 is associated with poor prognosis. A. Expressions of RIPK4 were analyzed in different tissues from four CRC cohorts. Gene expression data (GSE41258, GSE9348, GSE20916, GSE28702) were downloaded from the GEO database (*p<0.05, ** p<0.01, *** p<0.001, **** p<0.0001). B. Representative examples of RIPK4 immunohistochemical staining on normal tissue, primary CRC lesions, and liver metastases from the same patient from SYSUCC. C. Quantitative expression of RIPK4 in normal tissues, CRC, and liver metastases in SYSUCC. D. Kaplan Meier curves for overall survival, disease-specific survival, disease-free survival, and progression-free survival of CRC patients in the TCGA-COADREAD cohort. E. Kaplan Meier curves for overall survival and disease-free survival of CRC patients in the SYSUCC. F. RIPK4 protein content in 12 paired samples was detected by Western blot. G. RIPK4 protein content in samples from 2 patients with liver metastases. A, C. The mean ± s.d., unpaired two-tailed Student‘s t-test. Log-rank test was used for survival analysis. N, normal tissues; T, tumor tissues; LM or M, liver metastases. L, low expression; H, high expression. HR, Hazard Ratio. ns, not significant.

### RIPK4 promotes CRC anoikis resistance and metastasis

To explore the role of RIPK4 in CRC, we first detected the expression of RIPK4 in different CRC cell lines. RIPK4 was highly expressed in SW620 and LoVo cell lines from metastatic CRC (**Fig. 2A and Fig. 2B**). In a previous study, we found that RIPK4 levels increased during suspension (**Supplementary Fig. S1D and Supplementary Fig. S2A**). To further elucidate the relationship between elevated RIPK4 levels and suspension time, we measured RIPK4 mRNA and protein levels at different time points. Interestingly, the mRNA levels of RIPK4 decreased slightly on the first and second days of suspension, but increased significantly on the third days (**Fig. 2C**). In contrast, the protein level of RIPK4 increased continuously with increasing of suspension time (**Fig. 2D**). Next, we used sgRNA to construct stable lines of RIPK4 knockout CRC cells. As shown in the **Fig. 2E**, ko1 and ko2 showed good knockout efficiencies. After knockout of RIPK4, CRC cells showed reduced proliferation and mobility (**Supplementary Fig. S2B and Supplementary Fig. S2C**). Apoptosis increased in both adherent and suspended states (**Fig. 2F and Fig. 2I**). An in vivo lung metastasis model was established by injecting CRC cells into the tail vein of nude mice. We found that RIPK4 knockout significantly reduced the number of lung metastases, and the difference was statistically significant (**Fig. 2H**). The number and area of lung metastases in the knockout group were smaller than those in the control group, and the tumors were more prone to necrosis (**Fig. 2G**). Therefore, the results of this study confirmed that RIPK4 can resist anoikis and promote CRC cell metastasis.

**Fig. 2.**
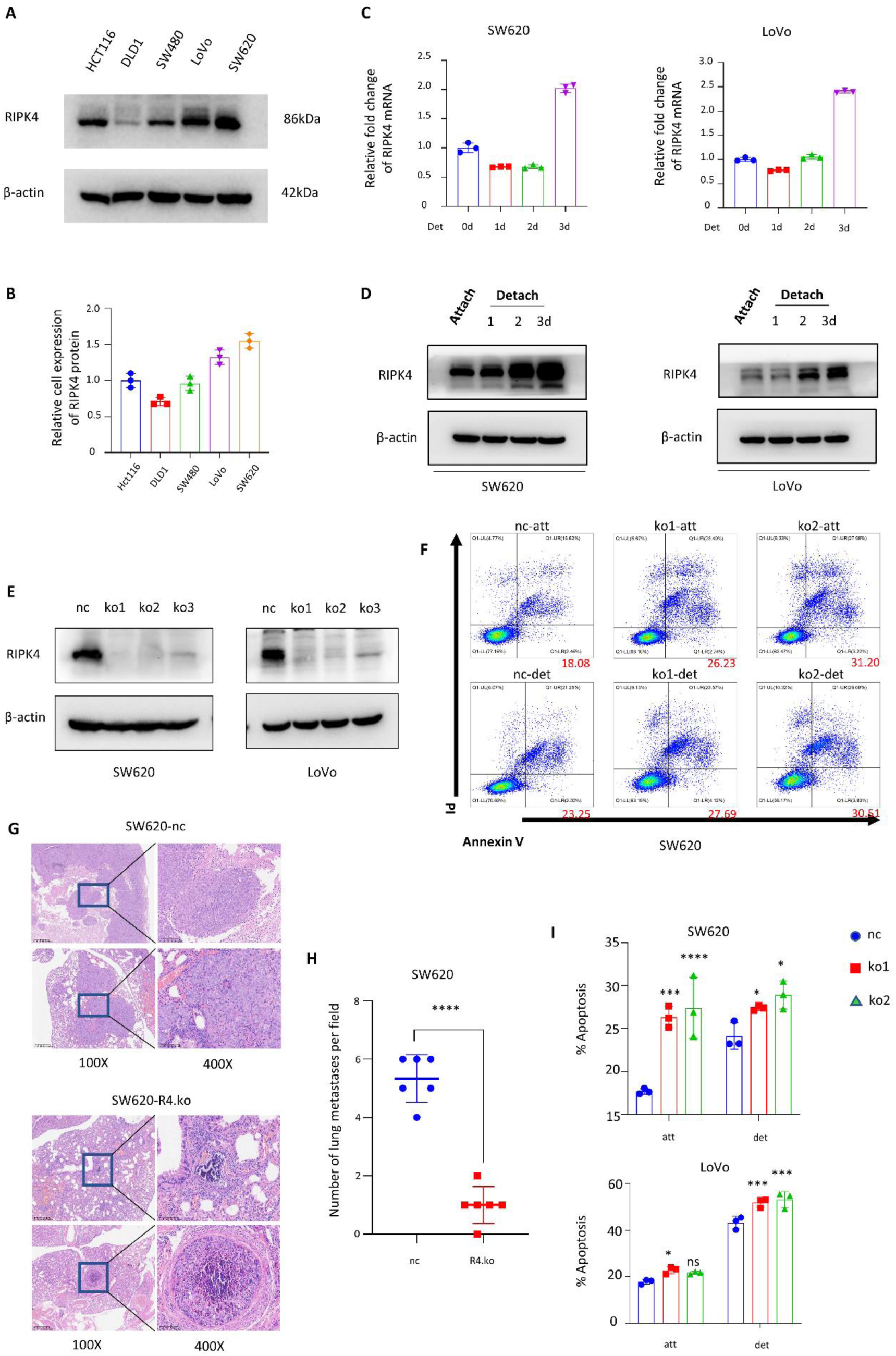
RIPK4 promotes CRC anoikis resistance and metastasis. A. Immunoblot analysis of WCL derived from different CRC cell lines. B. Quantitative analysis of RIPK4 derived from different CRC cell lines. C. RIPK4 mRNA changes D. RIPK4 protein changes in SW620 and LoVo cells with increasing suspension time. E. Western blotting of RIPK4 protein in SW620 and LoVo cells transfected with RIPK4 specific sgRNAs (ko1, ko2, ko3) or control sgRNAs. F. The proportion of SW620 apoptosis and anokis in specific culture conditions. G. Representative images of lung metastases in nude mice. H. Quantitative statistics of the number of lung metastases (n=6). Data are the mean ± s.d.; n = 6 biologically independent mice, unpaired two-tailed Student‘s t-test. I. Quantitative analysis of the proportion of SW620 and LoVo apoptosis and anokis under specific culture conditions. B, C, I. The mean ± s.d. for n = 3 independent experiments is shown. Statistical analysis was performed using an unpaired two-tailed Student‘s t-test. nc, normal control; ko, knockout; att, attached culture condition; det, detached culture condition.

### High expression of RIPK4 promotes tumor cells PANoptosis resistance

An increasing number of cell death models have been identified, including necroptosis, pyroptosis, and ferroptosis(*10, 41–43*). As described earlier, not only apoptosis but also other types of cell death occurred in suspended tumor cells. To verify the effect of RIPK4 on other cell death modes, we validated the cell death pathways involved in the RIPK4 protein using several inhibitors (Z-VAD-FMK, apoptosis inhibitor; Nec, necroptosis inhibitor; Fer, ferroptosis inhibitor; DMPD/DMLD, pyroptosis inhibitor; 3-me, autophagy inhibitor; dis, pyroptosis inhibitor). Unexpectedly, the inhibitor of the cell death pathway was unable to counteract the cell death effect caused by the knockout of RIPK4, either in the adherent state or in the suspended state. In the suspended state, the use of pyronecrosis and necrosis inhibitors may further promote tumor cell death (**Fig. 3A and Fig. 3B**). Similarly, we observed the fluorescence intensities of YP1 and PI at different suspension times. As shown in **Fig. 3C and Fig. 3D**, the fluorescence signals of YP1 and PI in RIPK4 knockout CRC cells increased significantly after 48 hours of suspension, and the differences were statistically significant (**24h and 72h in Supplementary Fig. S3**). After the knockout of RIPK4, the PI signal in CRC cells increased more significantly with increasing suspension time (**Fig. 3E and Fig. 3F**). Subsequently, we performed a soft agar colony formation assay on CRC cells to observe the ability of cell spheroids to survive in suspension. RIPK4 knockout CRC cells showed significantly reduced spheroidality in the half-suspended state and significantly reduced clone diameter (**Fig. 3G and Fig. 3H**). Thus, we found that RIPK4 promotes PANoptosis resistance in CRC cells rather than only apoptosis.

**Fig. 3.**
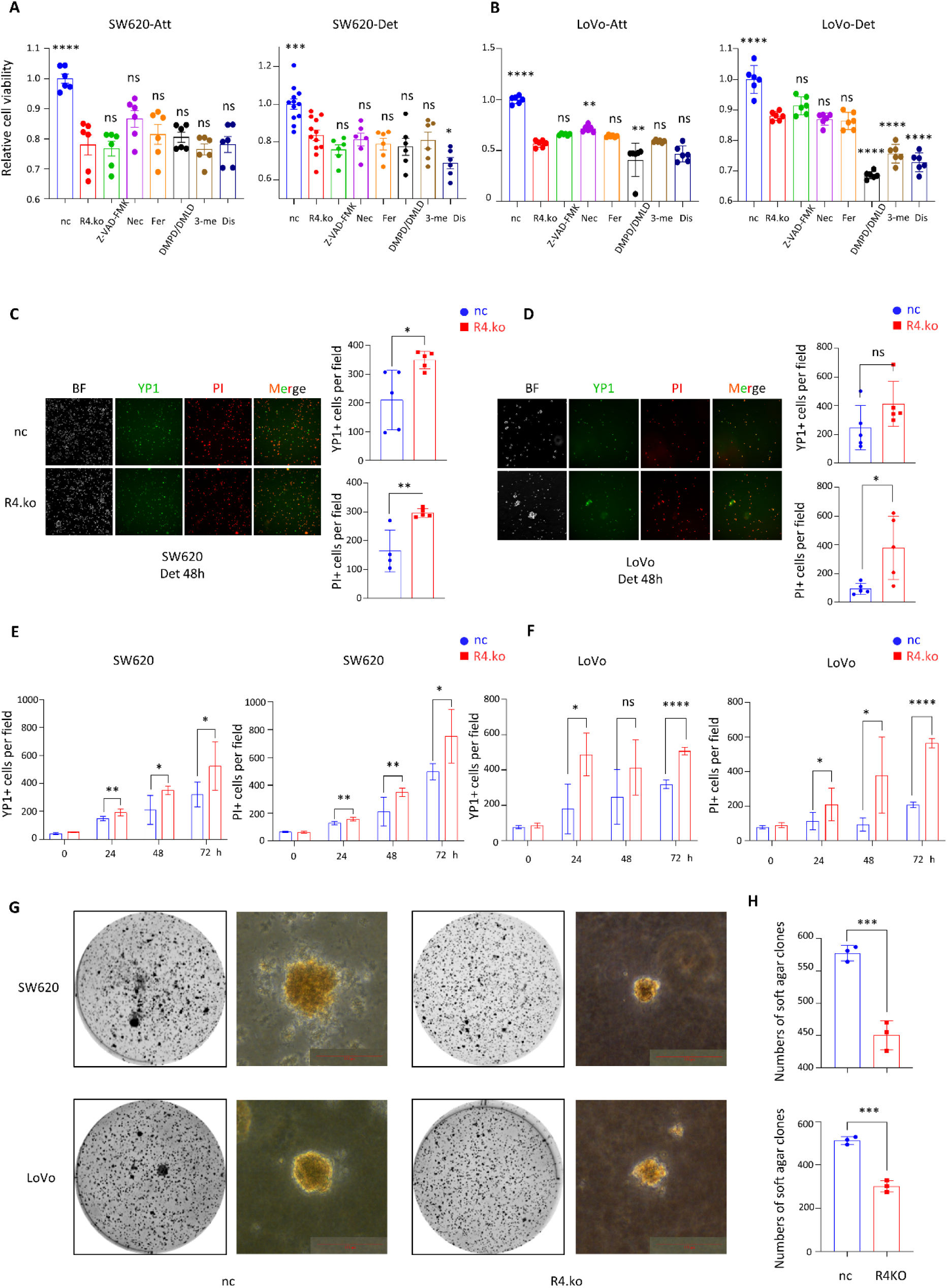
High expression of RIPK4 promotes tumor cells PANoptosis resistance. A-B. Cell viability of SW620 and LoVo cultured for 48 hours using cell death pathway inhibitors in attached or detached states. The dosage of inhibitors is described in the methodology section. C-D. YP1 and PI signals of SW620 and LoVo cultured in suspension for 48 hours. E-F. The relationship between YP1 and PI signals of SW620 and LoVo and changes in suspension time. G. Representative images of spheres under soft agar cloning and phase contrast microscopy. Scan bar = 100um. H. Quantitative data of cloned balls. The mean ± s.d. for n = 3 independent experiments, unpaired two-tailed Student‘s t-test. A-F. The mean ± s.d. for n = 6 independent experiments is shown. Statistical analysis was performed using an unpaired two-tailed Student‘s t-test. Z-VAD-FMK, apoptosis inhibitor; Nec, necroptosis inhibitor; Fer, ferroptosis inhibitor; DMPD/DMLD, pyroptosis inhibitor; 3-me, autophagy inhibitor; dis, pyroptosis inhibitor.

### RIPK4 promotes intracellular NADPH production and ROS clearance

Next, we aimed to elucidate the mechanism by which RIPK4 promotes CRC resistance to PANoptosis. Transcriptome sequencing analysis revealed that the RIPK4 was associated with NADPH production (**Fig. 4A**). It has been reported that NADPH in cells is mainly used to resist oxidative stress in cells, that is, scavenging reactive oxygen species (ROS)(*29, 44*). N-acetyl-L-cysteine (NAC) was a ROS inhibitor. In soft agar clones, NAC reversed the cell death caused by the knockout of RIPK4 (**Fig. 4B and Fig. 4C**). ROS levels in CRC cells were detected using DCFHDA probe. We found that the level of ROS in CRC cells increased after RIPK4 knockout (**Fig. 4D and Fig. 4E**). A decrease in NADPH levels was also observed in the RIPK4 knockout cells (**Fig. 4F and Fig. 4G**), consistent with our transcriptome sequencing results (**Fig. 4A**). Cell viability and NADPH levels were reversed by NAC (**Fig. 4H-K**). Therefore, it is possible that RIPK4 maintains the viability of tumor cells and induces PANoptosis resistance through the production of NADPH and the elimination of ROS. We also detected protein indicators of cell death related pathways.: Apoptosis caspase family; Necroptosis-associated RIPK1, RIPK3 and MLKL; LC3 and Beclin1 of autophagy and mitochondrial autophagy, and GSDM-family proteins involved in pyroptosis.

**Fig. 4.**
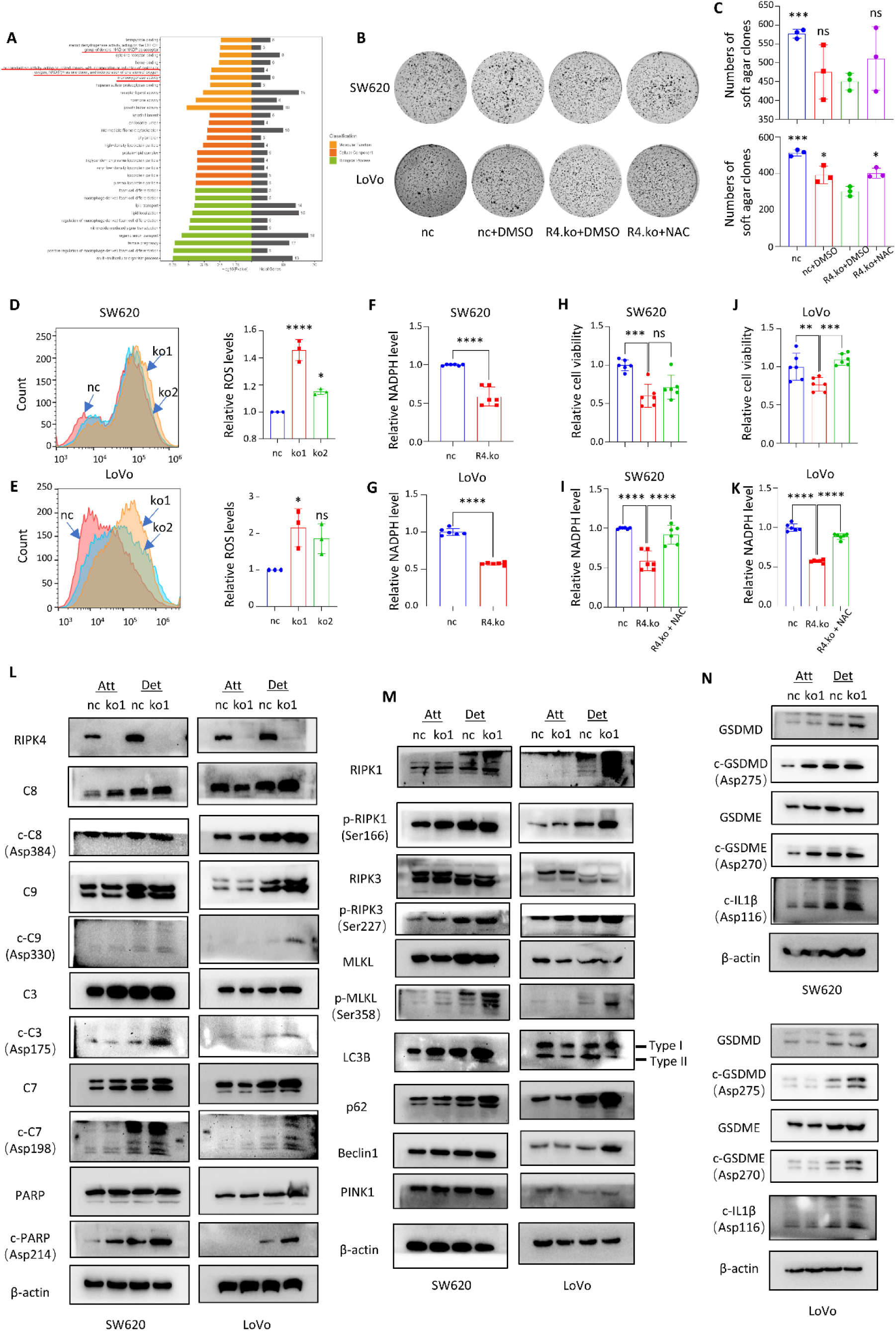
RIPK4 promotes intracellular NADPH production and ROS clearance. A. Protein functional signaling pathways recruited through RNA-seq by knocking out RIPK4. Pathway analysis showed that the change of RIPK4 expression caused the change of NADPH content in cells (the red line in the figure). B. Soft agar cloning under different treatment conditions. NAC, N-acetyl-L-cysteine, inhibitor of ROS. C. Quantitative counting of soft agar clones under different treatment conditions. RIPK4 knockout results in fewer clone spheres, which can be partially restored by NAC. D-E. Changes and quantitative analysis of intracellular ROS levels after knocking out RIPK4. Using DCFHDA probe for flow cytometry, the intracellular ROS level increased after RIPK4 knockout. F-G. Quantification of intracellular NADPH levels after knocking out RIPK4. H-I. Changes in intracellular NADPH after using NAC to clear ROS. J-K. Changes in cell viability after using NAC to clear ROS. L. Molecular changes in intracellular apoptotic pathways after knocking out RIPK4 in attach and detach states. M. Molecular changes in necroptosis and autophagy signaling pathways. N. Molecular changes in the cell pyroptosis pathway. A-E. The mean ± s.d. for n=3 independent experiments is shown. F-K. The mean ± s.d. for n=6 independent experiments is shown. The Western blot was repeated at least three times independently. Statistical analysis was performed using an unpaired two-tailed Student‘s t-test.

We further investigated how RIPK4 knockout affected these cell death pathways at the molecular level during suspension.We found that RIPK4 knockout significantly increased the cleavage forms of executioners Caspase-3, Caspase-7, and PARP in suspended CRC cells (**Fig. 4L**). Caspase-8 and Caspase-9, which act as upstream activators, showed significant increases in expression and cleavage during suspension after RIPK4 knockout (**Fig. 4L**). In suspended RIPK4 knockout CRC cells, significant pseudokinase mixed lineage kinase-like domain (MLKL) phosphorylation, a trigger of necroptosis, was observed (**Fig. 4M**). RIP1 and RIP3 are upstream activators of necroptosis. We observed a similar trend of increasing phosphorylation levels of RIPK1 and RIPK3 (**Fig. 4M**). At the same time, RIPK1 also had up-regulated expression and increased cleavage forms (**Fig. 4M**). We observed elevated expression of p62 and Beclin1 proteins in RIPK4 knockout suspension cells. The expression level of p62 protein was negatively correlated with the degree of autophagy(*45*). This was consistent with previous results, where the use of autophagy-related inhibitors led to further death of suspended tumor cells (**Fig. 3A-B**).

The downstream effector molecules of pyroptosis include Gasedermin D (GSDMD) and Gasedermin E (GSDME)(*46*). In RIPK4 knockout suspension cells, the expression of GSDMD and GSDME was up-regulated, and the cleavage forms were significantly increased (**Fig. 4N**). The same trend was observed in the cleavage of IL1β, which is an active cytokine produced during pyroptosis (**Fig. 4N**). RIPK4 gave CRC cells the property of being resistance to PANoptosis and can be used as a therapeutic target for tumor metastasis.

### RIPK4 interacts with MTHFD1

We explored the mechanism by which RIPk4 promotes NADPH production in CRC cells. The RIPk4 binding protein was stained with Coomassie blue and analyzed by mass spectrometry (**Fig. 5A**). Among the identified protein mixtures, the most abundant MTHFD1 protein was one of the key enzymes involved in the folate metabolism cycle (**Fig. 5B**). MTHFD1 exists mainly existed in the cytoplasm and catalyzes (6R)-5,10-methylene-5,6,7,8-tetrahydrofolate to produce (6R)-5,10-methyltetrahydrofolate and NADPH (**Supplementary Fig. S4A**)(*47–49*). We then validated the expression levels of MTHFD1 in suspended tumor cells, as shown in **Fig. 5C and Supplementary Fig. S4E**. MTHFD1 protein levels in the suspended tumor cells did not change (**Fig. 5C**). Therefore, we speculated that the effect of RIPK4 on the MTHFD1 protein is related to post-translational modifications. We used exogenous and endogenous co-ip experiments to verify the interaction between RIPk4 and MTHFD1 (**Fig. 5D and Fig. 5E**). Immunofluorescence analysis showed that they were located in the cytoplasm (**Fig. 5F**). In vitro GST-pulldown experiments showed that RIPK4 directly bound to MTHFD1 (**Fig. 5G**).

**Fig. 5.**
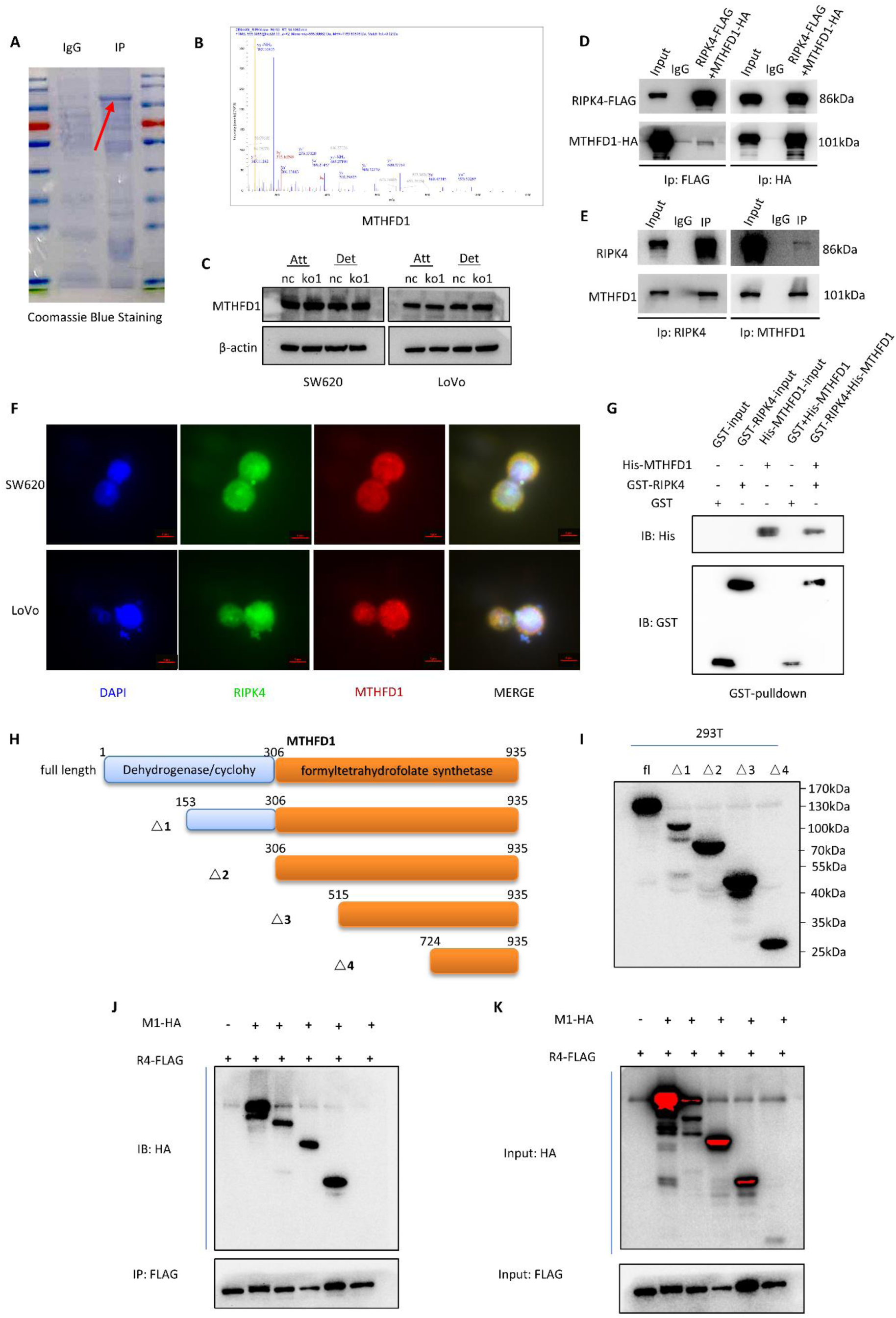
RIPK4 interacts with MTHFD1. A. Coomassie brilliant blue staining of RIPK4 binding protein. The red arrow shows the location of MTHFD1 protein. B. Mass spectrometry identification of MTHFD1 protein. C. The MTHFD1 protein levels did not change after suspension for 48 hours. D-E. Exogenous and endogenous co-ip binding experiments of RIPK4 and MTHFD1. E. Immunofluorescence showed co-localization of RIPK4 and MTHFD1 in the cytoplasm. Scan bar = 5um. G. GST-pulldown showed a direct binding relationship between RIPK4 and MTHFD1. H. Schematic diagram of MTHFD1 truncated plasmid construction. I. MTHFD1 truncated plasmid was validated as peptide by immunoblotting. J-K. Binding experiment between MTHFD1 truncated plasmid and RIPK4. The truncated MTHFD1-Δ4 could not bind to RIPK4, so it was inferred that the binding region of RIPK4 and MTHFD1 was between 515-724 amino acids of MTHFD1. C-K. Images are representative of n = 3 biologically independent experiments. IP, immunoprecipitation; fl, full length; M1, MTHFD1.

To clarify the specific binding regions of RIPK4 and MTHFD1, we constructed a truncated plasmid of MTHFD1 (**Structure diagram Fig. 5H**) and verified that the truncated plasmid could form peptides (**Fig. 5I**). In further truncated co-ip experiments, we found that MTHFD1 Δ4 lost its interaction with RIPk4, therefore we speculated that RIPk4 bound between MTHFD1 amino acids 515 to 724 (**IP, Fig. 5J and INPUT, Fig. 5K**).

### RIPK4 directly phosphorylates MTHFD1 545 threonine

Considering the characteristics of RIPK4 as a serine/threonine kinase, we performed quantitative co-ip experiments to detect the phosphorylation levels of total serine and threonine residues in MTHFD1. We observed that after RIPK4 knockdown, the total threonine phosphorylation level of MTHFD1 decreased, rather than the total serine phosphorylation level (**Fig. 6A**). In addition, we constructed RIPK4 inactivated plasmid RIPK4-I121N(*50*). RIPK4-I121N inactivated protein turned off its phosphorylation effect on MTHFD1 (**Fig. 6B**). This indirectly demonstrated the direct phosphorylation effect of RIPK4 on MTHFD1 rather than acting as a scaffold protein. In vitro phosphorylation kinase experiments directly demonstrated the effect of RIPK4 on MTHFD1 via phosphorylation modification (**Fig. 6C**). When we reintroduced the wild-type RIPK4 plasmid into the RIPK4 knockout cell line, the vitality and intracellular NADPH content of CRC cells was restored, whereas the inactivated RIPK4 plasmid did not have this effect (**Fig. 6D-G**). These results demonstrated the role of RIPK4 in the phosphorylation of MTHFD1 with threonine.

**Fig. 6.**
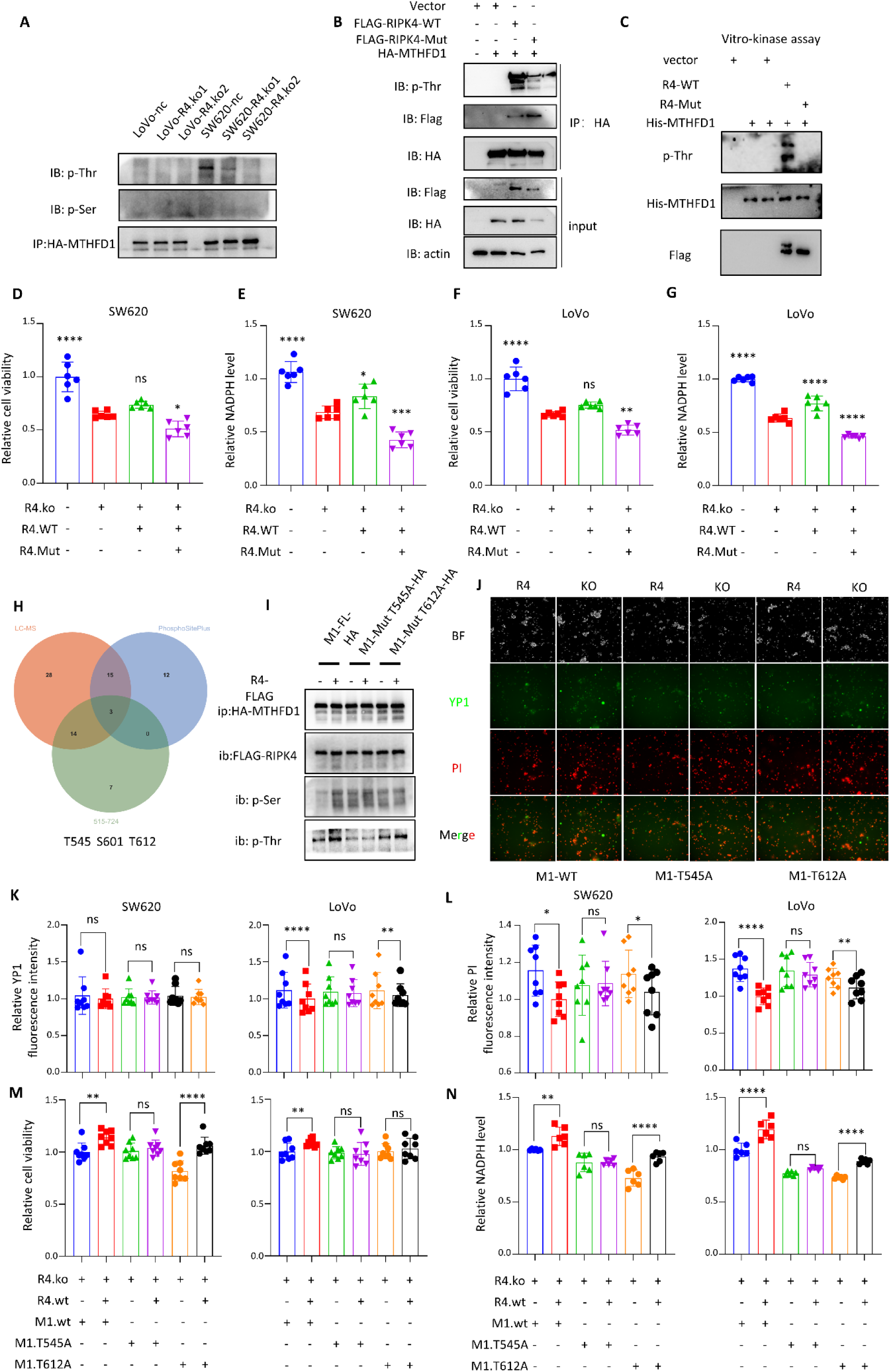
RIPK4 directly phosphorylates MTHFD1 545 threonine. A. The phosphorylation levels of threonine and serine residues in MTHFD1 after knocking out RIPK4 protein. After RIPK4 knockout, the level of threonine phosphorylation in MTHFD1 was significantly reduced, but not serine phosphorylation. B. The effect of RIPK4-WT or RIPK4-I121N on the threonine phosphorylation level of MTHFD1. RIPK4 phosphorylates MTHFD1 threonine residues, while RIPK4 protein inactivation mutation I121N loses the function of phosphorylating MTHFD1 protein. C. In vitro kinase experiments were conducted to verify the direct phosphorylation effect of RIPK4 on MTHFD1. D-G. The effect of RIPK4-WT/I121N on cell viability and NADPH. In RIPK4 knockout cells, the ectopic expression of RIPK4-WT/MUT protein, the intracellular NADPH level and cell viability can be saved by RIPK4-WT. H. The Venn diagram shows potential sites for the phosphorylation of MTHFD1 by RIPK4. The phosphorylation sites identified by mass spectrometry intersected with the phosphorylation sites in the public database (www.phosphosite.org). There were three phosphorylation sites in MTHFD1 515-724 amino acids. There were two threonine phosphorylation sites and one serine phosphorylation site. I. Quantitative co-ip analysis suggested that MTHFD1-T545 may be the phosphorylation site of RIPK4. J. Changes in YP1 and PI signaling after intracellular transfer of MTHFD1 point mutation plasmid. K-L. Quantitative analysis of changes in YP1 and PI signals after intracellular transfer of MTHFD1 point mutation plasmid. M-N. Quantitative analysis of changes in cell viability and NADPH after intracellular transfer of MTHFD1 point mutation plasmid. A-C, I. Images are representative of n = 3 biologically independent experiments. D-G, K-N. The mean ± s.d. for n = 6 independent experiments is shown. Statistical analysis was performed using an unpaired two-tailed Student‘s t-test. wt, wide type; mut, mutant type.

Previously, we identified RIPK4 binding between MTHFD1 amino acids 515 and 724 (**Fig. 5J and Fig. 5K**). To determine the specific binding sites of both, we performed mass spectrometry for phosphorylation site identification. Combining the MTHFD1 phosphorylation site we identified with the public database PhosphoSitePlus https://www.phosphosite.org/homeAction, three sites were found: T545, S601, and T612 (**Fig. 6H**, **Supplementary Fig. S4C and Supplementary Fig. S4D**). Considering that RIPK4 mainly phosphorylates threonine in MTHFD1, we constructed two point mutation plasmids, MTHFD1 T545A and T612A. We found that when MTHFD1 T545 was mutated, RIPk4 lost the phosphorylation of MTHFD1, whereas mutation T612 did not (**Fig. 6I**). We also performed YP1 and PI staining of tumor cells transfected with the MTHFD1 point mutation plasmid (**Fig. 6J-K**). When the T545 site was mutated, RIPK4 weakened its effect on cells through MTHFD1, in terms of both cell viability and intracellular NADPH (**Fig. 6M-N**). These results suggest that RIPK4 can directly phosphorylate threonine at the MTHFD1 545 site.

### RIPK4 functions with MTHFD1 dependence

To verify whether the function of RIPk4 is MTHFD1 dependent, we overexpressed RIPk4 in the cell, knocked down MTHFD1, performed both simultaneously, and detected the intracellular protein expression levels of both (**Supplementary Fig. S5A**). The Yp1 and PI signals suggested that knocking down MTHFD1 may restore the fluorescence changes caused by of RIPk4 overexpression (**Supplementary Fig. S5B-D**). Similar trends were observed for cell viability and intracellular NADPH levels (**Supplementary Fig. S5E-F**). Therefore, we determined that the function of RIPk4 was MTHFD1-dependent.

In further experiments, we constructed an MTHFD1 T545 site-specific phosphorylated antibody, and the dot blot test of the antibody is shown in **Supplementary Fig. S5G**. When RIPk4 was knocked out in the cells, p-MTHFD1 (545) decreased significantly (**Supplementary Fig. S5H**). P-MTHFD1 (545) was detected in the tissues of two CRC patients with liver metastases. In liver metastases, MTHFD1 phosphorylation was high expression (**Supplementary Fig. S5I**). High expression of phosphorylated MTHFD1 was also detected in 10 cases of CRC in primary tumors compared with that in matched normal tissues (**Supplementary Fig. S5J**). Higher levels of MTHFD1 phosphorylation were observed in the liver metastases (**Supplementary Fig. S5K**).

### The TP53 mutation is the key factor driving the high expression of RIPK4

In previous results, we found the highest expression levels of RIPK4 in SW620 and LoVo cells, both of which originated from metastatic CRC patients, with SW620 cells showing the most significant expression levels (**Fig. 2A-B**). The SW620 cell line has a hot spot mutation in TP53, R273H, which is comparable to LoVo. Based on this, we speculated that TP53 mutations might be related to the high expression of RIPK4. In the GSE41258 dataset containing TP53 status, a significant increase in expression of RIPK4 expression was found in CRC patients with TP53 mutations (**Fig. 7A**). ΔNp63 is a truncated transcript of TP63 and has been reported to be a transcription factor for RIPK4(*51–54*). We transferred the two most common R273 mutants of TP53 into the cell, R273H and R273C(*40*). As shown in **Fig. 7B**, when R273H and R273C mutations occurred in the cells, the mRNA levels of ΔNp63 and RIPK4 increased significantly. Similar results were observed at the protein level (**Fig. 7C**).To further clarify the interaction between the TP53 mutant protein and the ΔNp63 protein, the co-ip experiment showed that wild-type p53 protein could bind ΔNp63 protein and increase its ubiquitination level (**Fig. 7D**). However, the mutant p53 protein lost the function of binding with the ΔNp63 protein and reduced the ubiquitination level of the ΔNp63 protein (**Fig. 7D**).

**Fig. 7.**
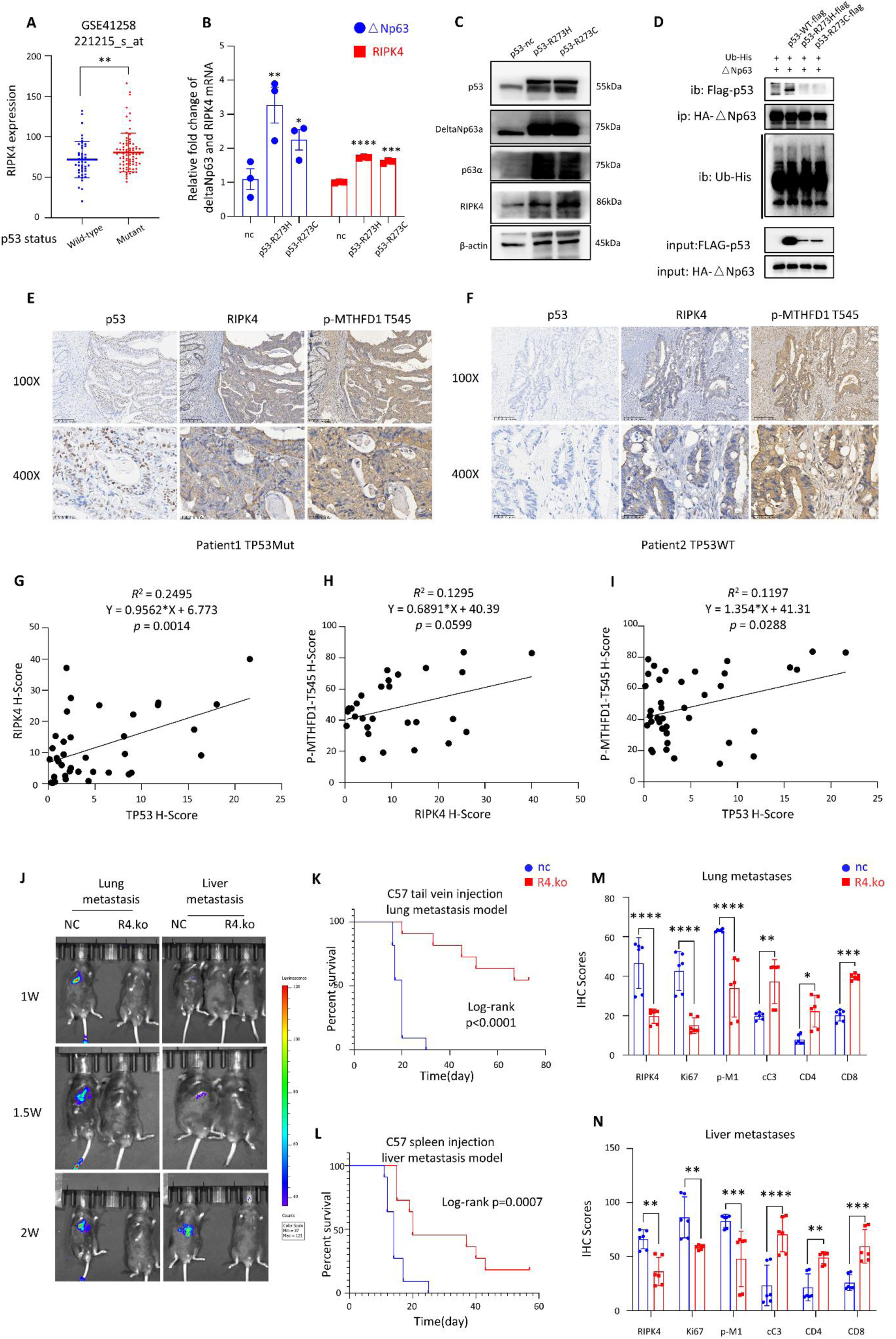
The TP53 mutation is the key factor driving the high expression of RIPK4. A. The expression levels of RIPK4 in the p53 WT/Mut states. The data was sourced from the GSE41258 cohort in the GEO database. B. After transferring the p53-R273H/C mutant plasmid into cells 48h, the mRNA of Δ Np63 and RIPK4 increased (n=3). C. After transferring the p53 R273H/C mutant plasmid into cells, the protein levels of Δ Np63 and RIPK4 increased. D. Quantitative co-ip experiments showed that the binding of p53 mutant protein to ΔNp63 was reduced, and its ubiquitination modification was reduced. E-F. Immunohistochemical staining of p53, RIPK4, and p-MTHFD1-T545 in CRC patients with TP53 mutation and wild-type. G-I. Quantitative and correlation analysis of p53, RIPK4, and p-MTHFD1-T545. J. Live imaging showed that in immunocompetent mice, knocking out RIPK4 significantly reduced the formation of liver and lung metastases. K-L. Survival curves of mice with liver and lung metastases (n=10). M-N. Immunohistochemical staining quantitative analysis of liver and lung tissues in mice, including RIPK4, Ki67, p-MTHFD1-T545, cleared caspase3, CD4, and CD8. In RIPK4 knockout mice, decreased MTHFD1 phosphorylation and Ki67, increased cell death cleaved caspase 3 and more lymphocyte infiltration were observed. A, B, M, N. The mean ± s.d., unpaired two-tailed Student‘s t-test. K, L. Log-rank test was used for survival analysis. C, D. The Western blot was repeated at least three times independently.

High expression of the RIPK4 protein and p-MTHFD1 T545 was also observed in patient tissues when the TP53 protein was mutated (**TP53 Mut, Fig. 7E; TP53 WT, Fig. 7F**). We observed that TP53 mutation status, RIPK4 and p-MTHFD1 T545 levels were positively correlated with each other in the tissues of patients with liver metastases (**Fig. 7G-I**). Therefore, mutant TP53 promotes RIPK4 transcription by reducing the inhibitory effect on Δ Np63, followed by RIPK4,which promotes intracellular NADPH generation and ROS clearance by phosphorylating MTHFD1, thereby promoting CRC PANoptosis resistance and distant metastasis.

We also observed the effect of RIPK4 using in vivo imaging of immune healthy C57 mice. MC38 cells with RIPK4 knocked out showed significantly reduced fluorescence signals, both in the lung and liver metastasis models (**Fig. 7J**). Mice bearing MC38 cells with RIPK4 knockout showed significant survival benefits (**Fig. 7K-L**). IHC staining showed more CD4+and CD8+T cells in RIPK4 knockout tumor tissues (**Fig. 7M-N and Supplementary Fig. S6A-B**).

## Discussion

Nearly half of patients with CRC experience distant metastases(*1*). This is one of the reasons for the poor prognosis and high mortality rate of CRC(*2, 3*). The discovery of novel metastatic mechanisms may provide new therapeutic targets and opportunities. In previous studies, the RIPK4 protein was shown to be differentially expressed in various cancer types, including ovarian cancer, melanoma, osteosarcoma, pancreatic cancer, bladder cancer, and nasopharyngeal carcinoma, and to participate in the EMT, Wnt, and NF-kB pathways to promote the occurrence and development of tumors(*14, 15, 21, 55–57*). In this study, we report that RIPK4 promotes tumor PANoptosis resistance and metastasis in CRC cells. To explore the mechanism of ROS generation and metastasis, we detected the expression of various proteins in suspended and attached CRC cells. Our data showed that the expression of RIPK4 was up-regulated in suspended cells, and the levels of RIPK1, RIPK3 and MLKL phosphorylation, and various types of cell death during suspension, including pyroptosis and necroptosis, were significantly increased in RIPK4-deficient CRC cells. Previous reports have showen that RIPK1 and RIPK3, which are critical for the formation of necrosomes, promote the process of necroptosis(*41*), (*58*). This indicated that RIPK4 exhibited completely different biological functions from RIPK1 and RIPK3, members of the same family(*59–61*).

Tumor cells require complete metastasis through the blood or lymphatic system, both of which are oxidative environments(*20, 62*). These diffuse tumor cells must overcome additional oxidative challenges(*22, 63–65*). Cancer cells require increased NADPH supplementation for redox homeostasis, lipid oxidation, and biomolecular synthesis, and several NADPH production pathways are enhanced in cancer cells(*29, 30*). The main intracellular source of cytoplasmic NADPH is the oxidative pentose phosphate pathway (oxPPP, ~ 30%), glutaminolysis flux through malic enzymes (~ 30%), and methylenetetrahydrofolate dehydrogenase–mediated folic metabolism (~ 40%)(*20*). Our data indicated that RIPK4 plays a key role in regulating redox homeostasis via promoting NADPH production.

Previous studies have demonstrated the functional properties of the protein kinases of the RIPK protein family: RIPK4 has been reported to phosphorylate plakophilin-1 (PKP1) during epidermal differentiation(*66*); RIPK4 also ensures epidermal differentiation and barrier function by phosphorylating the IRF6 signaling axis(*12*); During the process of necrosis, RIPK1 can be activated by self-phosphorylation, followed by the recruitment of RIPK3 to form necrosomes, which then phosphorylates MLKL, ultimately leading to necrosis(*59, 61*). To further study the mechanism by which RIPK4 regulates redox homeostasis, we immunoprecipitated the Flag-RIPK4 recombinant protein from the cell lysate of HEK293T and performed LC-MS/MS analysis. We identified MTHFD1 as a RIPK4-binding protein by the LC-MS/MS analysis. RIPK4 is a key upstream regulator of MTHFD1 that modulates MTHFD1 activity and function via direct phosphorylation of a specific threonine (Thr545).

MTHFD1, a key enzyme in the oxPPP pathway, is closely related to tumor disease progression and oxidative stress resistance(*22, 44, 67*). MTHFD1 is mainly located in the cytoplasm, whereas MTHFD1L and MTHFD2/2L are located in mitochondria(*68–70*). The MTHFD protein family plays an important role in tumor development. In non-small cell lung cancer (NSCLC), MTHFD1 silencing promotes cell apoptosis by inhibiting DNA methylation(*69*). Knockdown of MTHFD1L can reduce the proliferation of CRC, and in another study, knocking down MTHFD1L promoted the sensitivity of hepatocellular carcinoma to sorafenib(*68, 70*). In this study, we found no change in the expression of MTHFD1 in suspended tumor cells. However, RIPK4 dependent phosphorylation level of MTHFD1 was markedly up-regulated in detached cancer cells. Post-translational modifications (PTMs), including phosphorylation, ubiquitination, acetylation, and methylation, are well-known to change dynamically protein stability, localization, or catalytic activity. For example, phosphorylated NFS1 reduces the sensitivity of colorectal cancer to oxaliplatin chemotherapy by preventing PANoptosis(*7*). In recent studies, the increased methylation level of MTHFD1 enhanced its metabolic activity to produce NADPH, leading to anoikis resistance and distant organ metastasis(*71*). However, regulation of MTHFD1 function by phosphorylation has not been reported. Here, we reported that the phosphorylation of MTHFD1 plays an important role in regulating activity of MTHFD1.

In recent studies, RIPK2 has been identified as a key driver of immune evasion in cytotoxic T cell killing by reducing the level of MHC-I molecules on the cell surface through autophagy-lysosome degradation(*72*). The pyroptosis signal was significantly increased in tumor cells with RIPK4 deficiency. This may trigger the recruitment and infiltration of immune cells(*46, 59*). Thus, we observed that the deletion of RIPK4 in immune-competent mice led to significant survival benefits in mice with liver and lung metastases. Although we have not explored in depth the mechanism by which RIPK4 protein affects immunity, this will be one of the main focuses of our next work. Patients with TP53 mutation have a poor prognosis and early distant metastases(*33, 40*). Many studies have interpreted the biological behavior of malignancies caused by TP53 mutations in terms of GOF(*38–40, 73, 74*). In contrast to previous studies, we focused on post-translational modification of proteins and demonstrated the role of TP53 mutation/RIPK4/MTHFD1 axis in tumor PANoptosis resistance and metastasis. This clarifies a new mechanism for the poor clinical prognosis of TP53 mutations that may become a new therapeutic target.

In summary, upregulation of RIPK4 expression during suspension is crucial for tumor survival. We demonstrated that RIPK4 promoted tumor cell activity and metastasis by maintaining intracellular ROS homeostasis. Our findings also confirmed that TP53 mutations are a driver of RIPK4, suggesting that an in-depth study of this pathway may provide new therapeutic targets and strategies for metastatic CRC (**Fig. 8**).

**Fig. 8.**
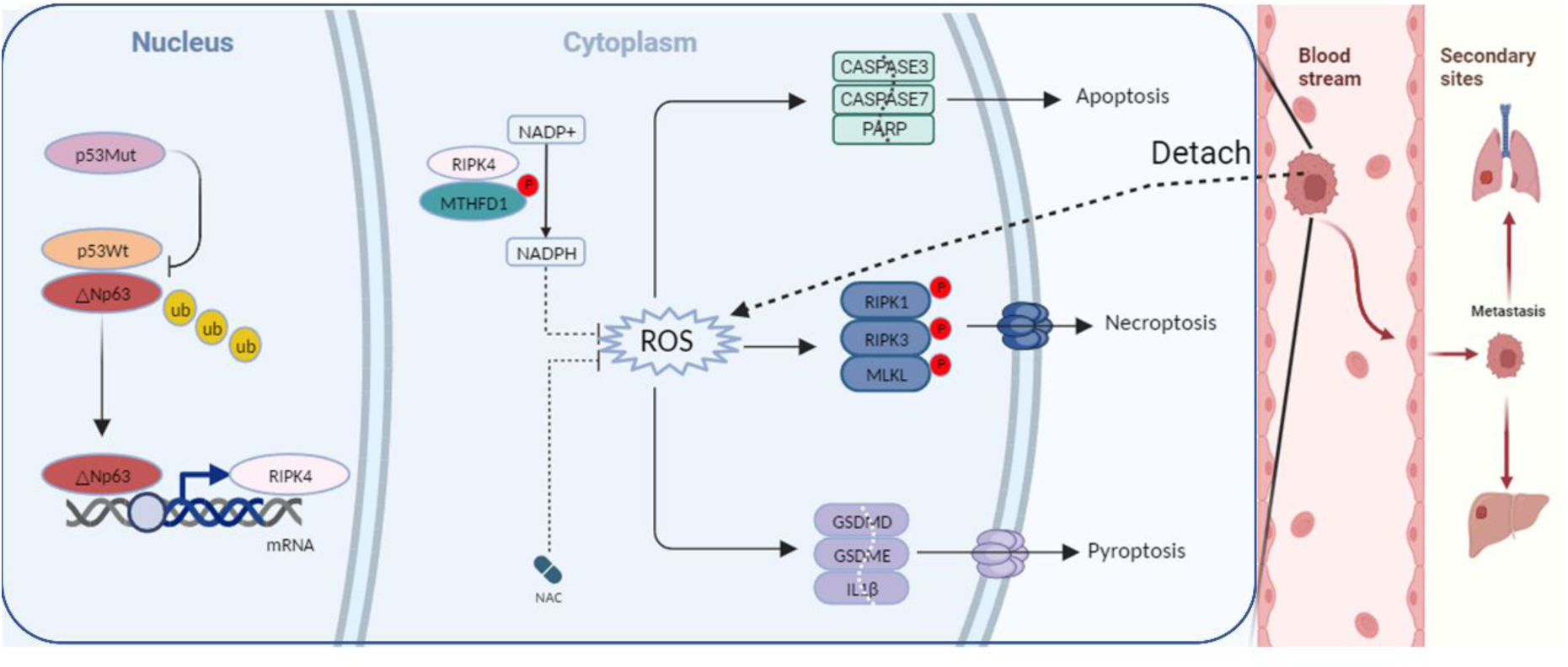
Schematic diagram of mechanism. **Nucleus:** The wild-type p53 protein binds to ΔNp63, increasing its ubiquitination modification. However, the mutant p53 protein loses this function, leading to the ΔNp63 binding to the promoter region of RIPK4 and promoting the transcription of RIPK4. **Cytoplasm:** Tumor cells in the circulation face oxidative stress, leading to an increase in intracellular ROS. The RIPK4 protein directly phosphorylates the MTHFD1 protein at Thr545 to generate more NADPH for ROS clearance. When the RIPK4 protein is knocked out, excessive accumulation of ROS in the cell triggered PANoptosis. Tumor cells, especially CRC with TP53 mutations, promote their survival in the circulatory system through the RIPK4/MTHFD1 axis.

## MATERIALS AND METHODS

### Cancer cell lines and inhibitors

The human colon cancer SW620 (CVCL_0547), huamn colon cancer LoVo (RRID:CVCL_0399), human HEK293T (RRID:CVCL_0063) and murine MC38 (CVCL_B288) cell lines were obtained from the American Type Culture Collection. SW620 and LoVo cells were cultured in 1640 medium containing 10% fetal bovine serum. The 293T (RRID: CVCL_0063) and MC38 (RRID: CVCL_B288) cells were grown in DMEM medium containing 10% fetal bovine serum. The incubation temperature was 37°C, and the incubator was moist with 20% O2 and 5% CO2. All cells were cultured in a 10 cm dish and sub-cultured into a six-well plate or a low-adhesion six-well plate for cell death, ROS and NADPH detection.

The cells were treated with reagents including apoptosis inhibitors Z-VAD-FMK (MedChenExpress, HY-16658B), necroptosis inhibitors Necrostatin-1 (MedChenExpress, HY-15760), ferroptosis inhibitors Ferrostatin-1 (MedChenExpress, HY-100579), pyroptosis inhibitors Ac-DMPD/DMLD-CMK and disulfiram (MedChenExpress, HY-16990 and HY-B0240), autophagy inhibitors 3-methyladenine (MedChenExpress, HY-19312), and ROS inhibitors Acetylcysteine (MedChenExpress, HY-B0215).

### Colorectal cancer tissues and IHC staining

The studies were performed with the approval from the Ethics Committee of Sun Yat-sen University Cancer Center. Fresh and formalin-fixed paraffin-embedded colorectal tumor samples were collected from patients diagnosed with CRC at Sun Yat-sen University Cancer Center using protocols approved by the institutional review board and following the Helsinki Declaration Code of Ethics. All patients received written informed consent and samples were identified prior to the trial. Tumor xenografts or tissue slides from surgical specimens were fixed in 10% neutral buffer formalin, embedded in paraffin, and cut into 4 mm sections. The samples were dewaxed and rehydrated before antigens were repaired with sodium citrate buffer. Tissue sections were stained with hematoxylin and eosin. For IHC staining, sections were incubated with 3% H2O2 for 15 minutes and blocked with 5% BSA at room temperature for 1 hour. The primary antibodies were incubated overnight and the cross sections were incubated with the corresponding secondary antibodies (ZSGB-BIO, PV-6000-55) at room temperature for 1 hour. All sections were stained with a DAB reagent kit (ZSGB-BIO, ZLI-9017) and the nuclei were re-stained with hematoxylin.

### Immunoblotting and immunoprecipitation

For immunoblotting, protein concentrations were measured after cell lysis using RIPA. Separate total protein (30ug) by SDS-PAGE and transfer to the PVDF membrane. Quantitative analysis of western blot was performed using Imagelab software. For both immunoprecipitation and immunocomprecipitation, NP40 lysate with phosphatase or protease inhibitor was used to lysate the cells. The protein supernatant was incubated with 2ug primary antibody at 4 °C overnight. On the second day, 30ul protein a/g agarose beads were added to the protein antibody mixture and incubated for 2 hours. Then the beads were washed five times with cold TBST, the supernatant was discarded, and the beads were heated to 100 °C with 50ul 1xsds for 10 minutes. Finally, western blot was performed.

Antibodies to the following were used: RIPK4 (CST, 12636, 1:1000), RIPK4 (ABclonal, A8495, for IHC 1:100), MTHFD1 (proteintech, 10794-1-AP, 1:1000), Beita-Actin (CST, 8457, 1:1000), DYKDDDDK-Tag (FLAG) (CST, 14793, 1:2000), HA-Tag (CST, 3724, 1:2000), GST-Tag (CST, 2625, 1:2000), His-Tag (CST, 2365, 1:2000), Phospho-Threonine (CST, 9386, 1:1000) (Jongsook Kim Kemper-University of Illinois at Urbana-Champaign Cat# phospho-Thr-SHP,RRID:AB_2889393), Phospho-Serine (Abcam, ab7851, 1;1000) (Abcam Cat# ab168254, RRID:AB_2572281), P53 (Abcam, ab32049, 1;1000) (Ollmann M; Cell. 2000Cat# p53, RRID:AB_2567126), Caspase-8 (CST, 4790, 1:1000), Cleaved Caspase-8 (Asp384) (CST, 9748, 1:1000), Caspase-9 (CST, 9502, 1:1000), Cleaved Caspase-9 (Asp330) (CST, 7237, 1:1000), Caspase-3 (CST, 14220, 1:1000), Cleaved Caspase-3 (Asp175) (CST, 9664, 1:1000), Caspase-7 (CST, 12827, 1:1000), Cleaved Caspase-7 (Asp198) (CST, 9491, 1:1000), PARP (CST, 9532, 1:1000), Cleaved PARP (Asp214) (CST, 5625, 1:1000), RIPK1 (proteintech, 17519-1-AP, 1:1000), Phospho-RIP (Ser166) (CST, 65746, 1:1000), RIPK3 (proteintech, 17563-1-AP, 1:1000), Phospho-RIP3 (Ser227) (CST, 93654, 1:1000), MLKL (proteintech, 21066-1-AP, 1:1000), Phospho-MLKL (Ser358) (CST, 91689, 1:1000), SQSTM1/p62 (CST, 23214, 1;1000), LC3B (CST, 83506, 1:1000), Beclin-1 (CST, 3495, 1:1000), PINK1 (CST, 6946, 1:1000), Gasdermin D (CST, 69469, 1:1000), Cleaved Gasdermin D (Asp275) (CST, 36425, 1:1000), Gasdermin E (CST, 19453, 1:1000), Cleaved Gasdermin E (Asp270) (CST, 55879, 1:1000), Cleaved-IL-1β (Asp116) (CST, 83186, 1:1000), CD4 (proteintech, 67786-1-Ig, for IHC 1:500), CD8 (proteintech, 29896-1-AP, for IHC 1:500), Ki67 (proteintech, 27309-1-AP, for IHC 1:2000).

### Cell-death assays

The cell death detection kit (Beyotime, C1075) was used and stained with YO-PRO-1 (YP1) and propidium iodide (PI). CRC cells were grown on 6-well plates or low-adhesion 6-well plates (Corning, 3471) and treated with YP1 and PI reagents. The labeled cells were observed under fluorescence microscope: YP1 positive cells (green) had apoptosis or necroptosis, and PI positive cells (red) had necroptosis, pyroptosis or ferroptosis. Quantitative analysis was performed using a fluorescence microplate reader.

### Cell viability assay

The cell viability was measured by alamarblue assay (ThermoFisher, DAL1025). Cells were seeded in 96 well plates with 10000 cells per well on a white background, with or without 6 mg/ml poly-HEMA (Sigma, P3932, RRID:SCR_000488) coating. The cell viability was measured at the specified time after incubation according to the manufacturer’s instructions.

### mRNA extraction, qPCR, and RNA sequencing

According to the manufacturer’s protocol, total RNA was extracted using Trizol (Invitrogen, 15596-026) method. Reverse transcription of total RNA was performed to generate cDNA. The polychromatic real-time PCR detection system (EZBioscience, A0012-R2) and SYBR Green real-time PCR Master Mix (EZBioscience, A0001-R1) were used for real-time PCR. The primer sequences used for real-time PCR analysis are listed below. ΔNP63α, Forward primer (5′-to3′), CTGGAAAACAATGCCCAGAC, Reverse primer (5′-to3′), GGGTGATGGAGAGAGAGCAT; TAp63, Forward primer (5′-to3′), TACTGCCCTGACCCTTACATC, Reverse primer (5′-to3′), TCTTGTTTGTCGCACCATCTT; RIPK4, Forward Primer (5′-to3′), AAACGGAATCAAGGGGACAGTT, Reverse Primer (5′-to3′), GCTTATCTGCCGTCGAACCA; Βeita-actin, Forward Primer (5′-to3′), GGACTTCGAGCAAGAGATGG, Reverse Primer (5′-to3′), AGCACTGTGTTGGCGTACAG.

Total RNA was isolated from cells/tissues using the Magzol Reagent (Magen, China) according to the manufacturer’s protocol, The quantity and integrity of RNA yield was assessed by using the K5500 (BeijingKaiao, China) and the Agilent 2200 TapeStation (Agilent Technologies, USA) separately. Briefly, the mRNA was enriched by oligodT according to instructions of NEBNext® Poly(A) mRNA Magnetic Isolation Module (NEB,USA). That was followed by a break-up to approximately 200bp. Subsequently, theRNA fragments were subjected to first- and second-strand cDNA synthesis following adaptor ligation and enrichment with low-cycle as described by. ofNEBNext® Ultra™ RNA Library Prep Kit for Illumina. Valuation of purified library products using the Agilent 2200 TapeStation and Qubit (Thermo Fisher Scientific, USA). The libraries were sequences by Illumina (Illumina, USA) with paired-end 150bp at Ribobio Co. Ltd (Ribobio, China). In order to identify the Gene Ontology (GO) annotations and pathways in which differentially expressed genes were enriched, GO term and Kyoto Encyclopedia of Genes and Genomes (KEGG, RRID:SCR_012773) pathway enrichment analyses were performed using the “clusterProfiler (RRID:SCR_016884)” package in RBioconductor /KOBAS3.0 software. In the present study, the clusterProfiler package in R Bioconductor/KOBAS3.0 was used to identify and visualize GO terms and KEGG pathways enriched by all differentially expressed genes.

### CRISPR-Cas9 mediated gene knockout

We used CRISPR-Cas9 technology to knock out RIPK4. Each sgRNA was cloned into the empty backbone of Lenti CRISPR v2. The following sgRNA sequences were used: 5’-CGTGAACTCGCCCGCGTCGA-3’, 5’-GTTCCGAATCATCCACGAGA-3’ and 5’-CTGCCGCGAACCTGTCGGCC-3’. The plasmids containing sgRNA sequence were transfected into HEK293T cells with psPAX2 (RRID: Addgene_12260) packaging plasmid and pMD2.G VSV-G envelope-expressing plasmid. After 72 hours, the virus was collected and infected with virus and 1ug/ml polybrene in SW620, LoVo and MC38 cells. Selected with puromycin (2ug/ml, Invivogen) for 1 week.

### Determination of ROS and NADPH

ROS detection was performed using a reactive oxygen species specification assay kit (Beyotime, S0033). Remove the cell culture medium and add the appropriate volume of diluted DCFH-DA. Incubate in 37°C cell incubator for 30 minutes. The cells were washed three times with serum-free cell culture medium to fully remove DCFH-DA that did not enter the cells. Cells were collected and detected by flow cytometry.

The NADPH test kit (Beyotime, S0179) was used to detect NADPH levels in the cells. Remove the cell culture medium, add the NADPH extract, and blow gently. 60°C water bath for 30 minutes to decompose NADP+. Add the G6PDH working solution and mix well. Incubate at 37°C in dark for 10 minutes. Add chromogenic solution and incubate at 37°C for 10-20 minutes in dark. The absorption was measured at 450nm.

### Soft agar colony formation assay

Prepare a 5% sterilized agar solution. After mixing with the medium, the lower layer of glue was laid in a 6-well plate with a concentration of 0.5%. After the lower layer of glue solidified, the upper layer of glue containing CRC cells (2000 cells per well) was laid with a concentration of 0.35%. After the upper gel has solidified, 200 UL of complete medium is added to each well to prevent the agar gel from drying out. 100ul medium was supplemented every 3 days. After 3 weeks of culture, the size and morphology of the CRC clone spheres were observed under a microscope. Clone balls were counted after 24 hours of incubation with MTT solution and were stained purple-black. Take pictures and count them with ImageJ (RRID:SCR_003070) software.

### Plasmid constructs

pcDNA3.1-RIPK4-WT-FLAG, pcDNA3.1-RIPK4-I121N-FALG, pcDNA3.1-TP53-WT-FLAG, pcDNA3.1-TP53-R273H-FLAG and pcDNA3.1-TP53-R273C-FLAG were cloned into mammalian expression pCDNA3.1-FLAG-Amp vectors. pcDNA3.1-MTHFD1-HA, pcDNA3.1-MTHFD1-T545A-HA, pcDNA3.1-MTHFD1-T6412A-HA, pcDNA3.1-MTHFD1-Δ 1-HA, pcDNA3.1-MTHFD1-Δ 2-HA, pcDNA3.1-MTHFD1-Δ3-HA, pcDNA3.1-MTHFD1-Δ4-HA and pcDNA3.1-ΔNP63-HA were cloned into mammalian expression pCDNA3.1-HA-Amp vectors.

### GST-pulldown assay

The recombinant proteins GST-RIPK4 and His-MTHFD1 were produced by transforming BL21 (DE3) Escherichia coli strain. GST-RIPK4 protein was incubated with GSH magnetic bead suspension at 4°C overnight. The next day, it was cleaned with a cold PBST on a magnetic frame for 5 epochs. Then His-MTHFD1 protein solution was added to GST-RIPK4 protein solution and incubated overnight at 4°C. On the third day, the precooled PBST was again washed on the magnetic frame 5 times, the elution buffer was added, and denaturated and eluted in a boiling water bath for 10 min. Finally, the reduced protein loading buffer was added and denatured for 10 minutes in a boiling water bath. Westernblot was used to detect the target protein.

### In vitro IP-kinase assay

Transfer RIPK4-WT-FLAG or RIPK4-I121N-FLAG mutant inactivated plasmids into HEK293T cells. After 48 hours, the cells were lysed with RIPA lysate and the supernatant was obtained after centrifugation. The FLAG magnetic beads were then incubated overnight and immunoblotted to detect IP proteins (**Supplementary Fig. S4F**). Expression of the recombinant protein pet28a-MTHFD1-6xHis using a prokaryotic vector system. Incubate RIPK4-WT-FLAG or RIPK4-I121N-FLAG kinases with MTHFD1 protein in 1X kinase buffer (Cell Signaling Technology, 9802), supplemented by 200uM cold ATP (Cell Signaling Technology, 9804) at 37 °C for 30 minutes. The kinase assay was terminated with a 20ul 3x SDS sample buffer. His antibody (Cell Signaling Technology, 12698) was used to detect the MTHFD1 protein, Flag antibody (Cell Signaling Technology, 14793) was used to detect the RIPK4 kinase, and threonine phosphorylation antibody (Cell Signaling Technology, 9386) was used to detect the phosphorylation level of the MTHFD1 protein.

### Animal experiments

This study complies with all relevant ethical regulations. All procedures and experimental protocols involving mice were approved by the institutional animal care and use Committee (IACUC) of Sun Yat-sen University Cancer Center. Female BALB/c nude and C57 mice were purchased from Charles River (Beijing), aged 4-6 weeks. All mice were raised at the Animal Experiment Center of Sun Yat-sen University under no specific pathogen conditions. To investigate the effect of RIPK4 on CRC metastasis, we injected 10E6 SW620-NC or RIPK4KO cells into BALB/c nude mice through the tail vein, respectively. After 8 weeks, the mice were sacrificed and lung tissue was obtained, and the number of lung metastases was observed under paraffin. Subsequently, in C57 mice with sound immune function, 5*10^5^ MC38-NC-Luciferase or MC38-RIPK4ko-Luciferase cells were injected into the tail vein or spleen capsule of mice. Live imaging fluorescence signals were observed every 3 days and D-Luciferin potassium salts were injected into the abdominal cavity. Mice survival times were recorded and all mice were sacrificed after eight weeks to obtain liver or lung tissue embedded in paraffin. Liver and lung metastases were observed under a paraffin cross-section microscope.

### Mass spectrometry

For in-gel tryptic digestion, gel pieces were destained in 50 mM NH4HCO3 in 50% acetonitrile (v/v) until clear. Gel pieces were dehydrated with 100 μl of 100% acetonitrile for 5 min, the liquid removed, and the gel pieces rehydrated in 10 mM dithiothreitol and incubated at 56 °C for 60 min. Gel pieces were again dehydrated in 100% acetonitrile, liquid was removed and gel pieces were rehydrated with 55 mM iodoacetamide. Samples were incubated at room temperature, in the dark for 45 min. Gel pieces were washed with 50 mM NH4HCO3 and dehydrated with 100% acetonitrile. Gel pieces were rehydrated with 10 ng/μl trypsin resuspended in 50 mM NH4HCO3 on ice for 1 h. Excess liquid was removed and gel pieces were digested with trypsin at 37 °C overnight. Peptides were extracted with 50% acetonitrile/5% formic acid, followed by 100% acetonitrile. Peptides were dried to completion and resuspended in 2% acetonitrile/0.1% formic acid. The tryptic peptides were dissolved in 0.1% formic acid (solvent A), directly loaded onto a home-made reversed-phase analytical column (15-cm length, 75 μm i.d.). The gradient was comprised of an increase from 6% to 23% solvent B (0.1% formic acid in 98% acetonitrile) over 16 min, 23% to 35% in 8 min and climbing to 80% in 3 min then holding at 80% for the last 3 min, all at a constant flow rate of 400 nl/min on an EASY-nLC 1000 UPLC system. The peptides were subjected to NSI source followed by tandem mass spectrometry (MS/MS) in Q ExactiveTM Plus (Thermo) coupled online to the UPLC. The electrospray voltage applied was 2.0 kV. The m/z scan range was 350 to 1800 for full scan, and intact peptides were detected in the Orbitrap at a resolution of 70,000. Peptides were then selected for MS/MS using NCE setting as 28 and the fragments were detected in the Orbitrap at a resolution of 17,500. A data-dependent procedure that alternated between one MS scan followed by 20 MS/MS scans with 15.0s dynamic exclusion. Automatic gain control (AGC) was set at 5E4.The resulting MS/MS data were processed using Proteome Discoverer (RRID:SCR_014477) 1.3. Tandem mass spectra were searched against SwissProt database. Trypsin/P (or other enzymes if any) was specified as cleavage enzyme allowing up to 2 missing cleavages. Mass error was set to 10 ppm for precursor ions and 0.02 Da for fragment ions. Carbamidomethyl on Cys were specified as fixed modification and oxidation on Met was specified as variable modification. Peptide confidence was set at high, and peptide ion score was set > 20.

### Peptide synthesis and Antibody purification

Two modified polypeptides and one unmodified polypeptide were designed and synthesized according to the sequence information by Jingjie PTM BioLab (Hangzhou) Co. Inc.. The sequences are as follows: MTHFD1-T545-Phospho1: CDRFLRKI-(phospho)T-IGQ, MTHFD1-T545-Phospho2: CTNDRFLRKI-(phospho)T-IGQ, MTHFD1-T545-WT: CTNDRFLRKITIGQ. Mass spectrometry was used to verify the purity of the synthetic polypeptides. The two modified polypeptides were conjugated with KLH for rabbit immunization. Each rabbit was immunized four times. On the 70th day, 30 mL of whole blood was collected and centrifuged to collect the supernatant for serum screening, including ELISA and dot blot. Positive serum was then selected for purification. The antibodies were validated using ELISA, Dot blot and Western Blot.

### ELISA and Dot blot assay

ELISA. The antigen was diluted with coating solution and 50 ug/well was added to the ELISA plate in sequence, followed by incubation at 4°C overnight or 37°C for 2 hours. The coated plate was washed three times with 1×TBST and then blocked with 1% BSA. The plate was incubated at 37°C for 1 hour before being washed for 1-3 times. The primary antibody was diluted 3-fold starting from 1:1,000 and added to the ELISA plate, which was then incubated at 37°C for 1.5 hours and washed for 1-3 times. The secondary antibody was diluted to 1:10,000 with 1% BSA and added to the plate, which was incubated at room temperature or 37°C for 45 minutes and washed 1-3 times. After adding TMB chromogenic solution for 5-10 minutes, the reaction was terminated with 1M sulfuric acid, and the data was read with a microplate reader.

Dot blot. Uncrosslinked antigenic peptides were spotted onto PVDF membranes in a gradient of 1 ng, 4 ng, 16 ng, and 64 ng. After the membrane surface was dried, the membrane was blocked at room temperature for 60 minutes and then washed with 1×TBST for 10 minutes. The primary antibody was diluted with 2.5-5% nonfat dry milk and added to the membrane for incubation at room temperature for 2 hours. Membrane was washed three times with 1×TBST for 5-10 minutes each time. Secondary antibody was applied at a dilution of 1:10,000 and incubated at room temperature for 45 minutes to 1 hour. Finally, the membrane was washed three times in 1×TBST (5-10 minutes per wash) and developed.

### Quantification and statistical analysis

All Western blotting and immunoprecipitation experiments were repeated independently at least three times with similar results. Other data (if noted) are expressed as mean ± standard deviation. Or the standard error of the mean (S.E.M.). A two-way ANOVA was used for the multi-group comparison and an unpaired two-tailed t-test was used for the two-group comparison. Significant statistical differences between groups were indicated as, ***\*P < 0.05, **P < 0.01, ***P < 0.001, ****P < 0.0001.*** GraphPad Prism 8.0 and Microsoft Excel (RRID:SCR_016137) 16.0 were used for statistical analysis and graphics.

## List of Supplementary Materials

Materials and Methods

Fig. S1 to Sx for multiple supplementary figures

## Author contributions

Z.-L.H.and L.Y. designed and conceptualized this project. L.Y., S.Z., and Y.-B.X. conducted most of the experiments. L.Y., Z.-J.Z. and X.-M.C. were responsible for animal experiments. C.Z., Q.-J.O. and Y.-J.F. analyzed the sequencing data. J.-H.P. and J.-Z.L. analyzed the clinical data. L.Y., B.L., Z.-Z.P. and Z.-L.H. wrote the manuscript. All co-authors have seen and approved the manuscript.

## Data availability

The MS data generated in this study has been saved in iProX/ProteomeXchange with login number IPX0007657000/PXD047592. The RNA sequencing dataset has been stored in the Gene Expression Omnibus with login number GSE252264. All other data supporting the findings of this study are available from the corresponding author upon reasonable request. All data are available in the main text or the supplementary materials.

---

**Supplementary Fig. S1.**
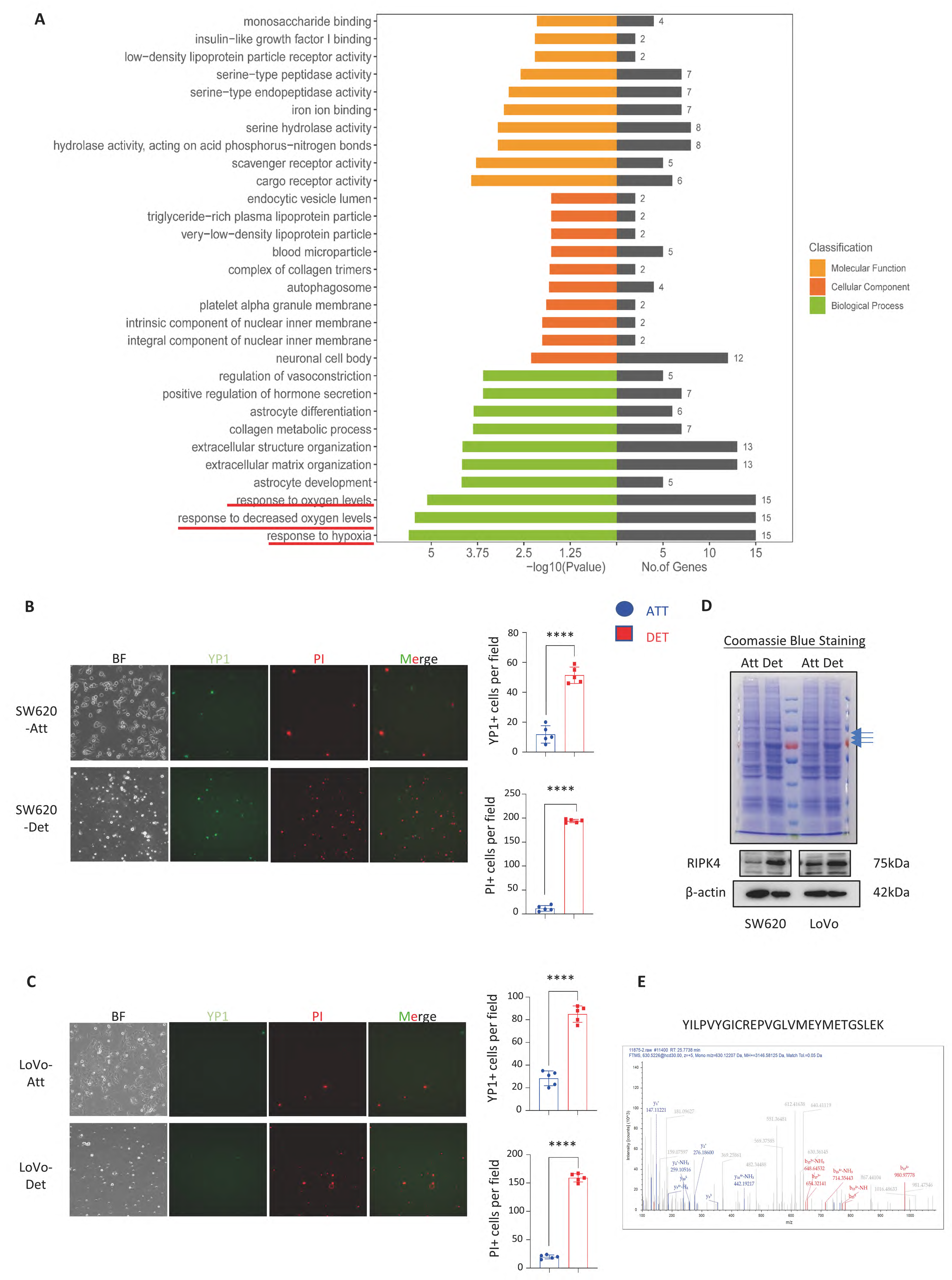
Tumor cell death during suspension. A. RNA seq of tumor cells in attach and detach states. Pathway analysis revealed hypoxia and oxidative stress within tumor cells (Red underlined part). B-C. The changes in YP1 and PI signals within tumor cells in attach and detach states suggested multiple cell death modes that tumor cells face during suspension. The signal intensities of YP1 and PI in suspension cells were significantly up-regulated, especially the PI signal. D. Coomassie bright blue staining showed protein changes within cells in a suspended state. E. Identification spectrum of RIPK4 protein by mass spectrometry. B, C. The mean ± s.d., n=6, unpaired two-tailed Student‘s t-test.

**Supplementary Fig. S2.**
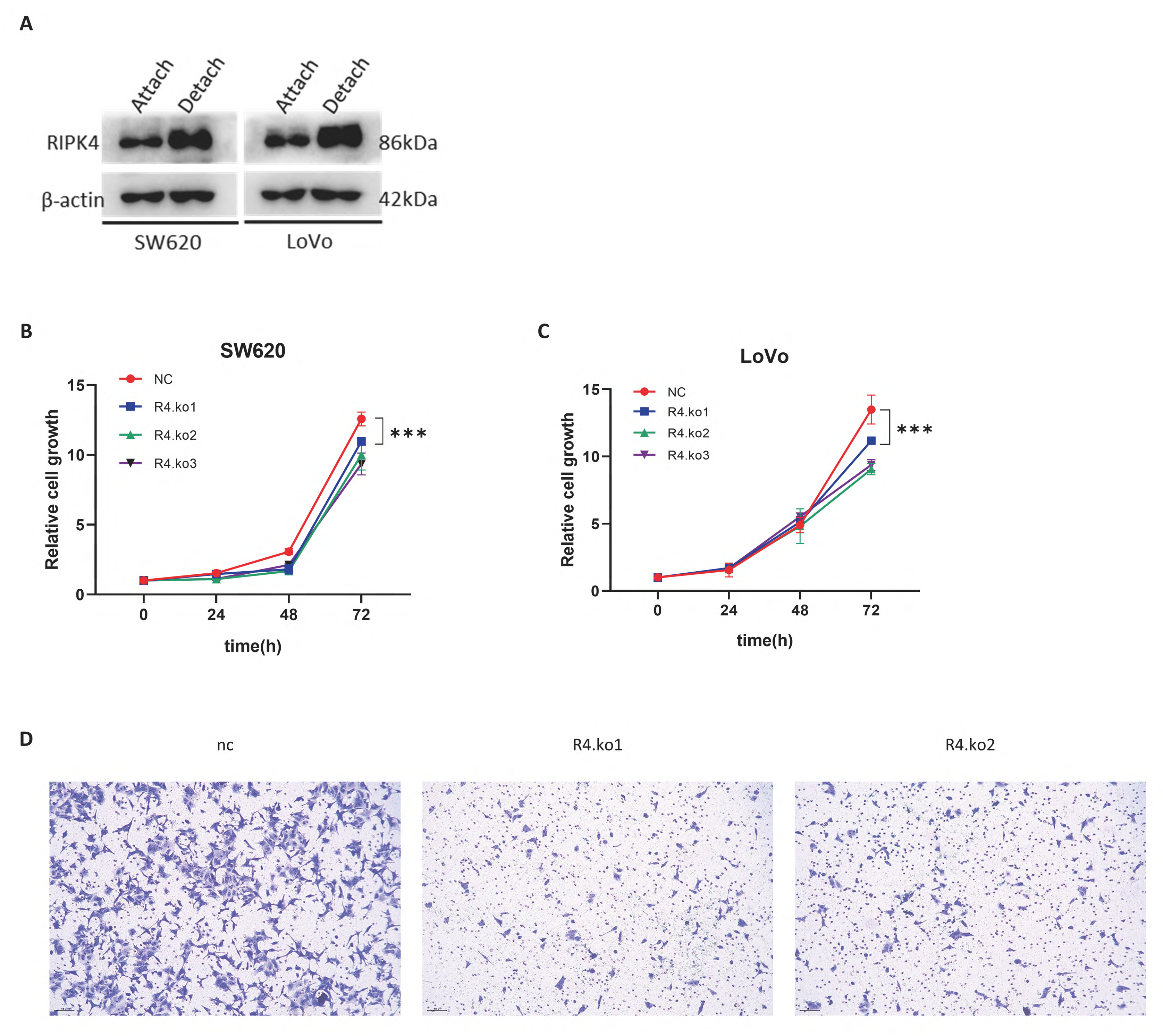
RIPK4 expression increased during cell suspension, and knocking out RIPK4 reduced the proliferation rate and migration ability of CRC cells. A. Immunoblotting showed an increase in the expression level of RIPK4 protein during the suspension process. B-C. The rate of cell proliferation decreased after knocking out RIPK4. D. After knocking out RIPK4, the migration ability of tumor cells weakened. The Western blot was repeated at least three times independently. B, C. The mean ± s.d., n=6, unpaired two-tailed Student‘s t-test.

**Supplementary Fig. S3.**
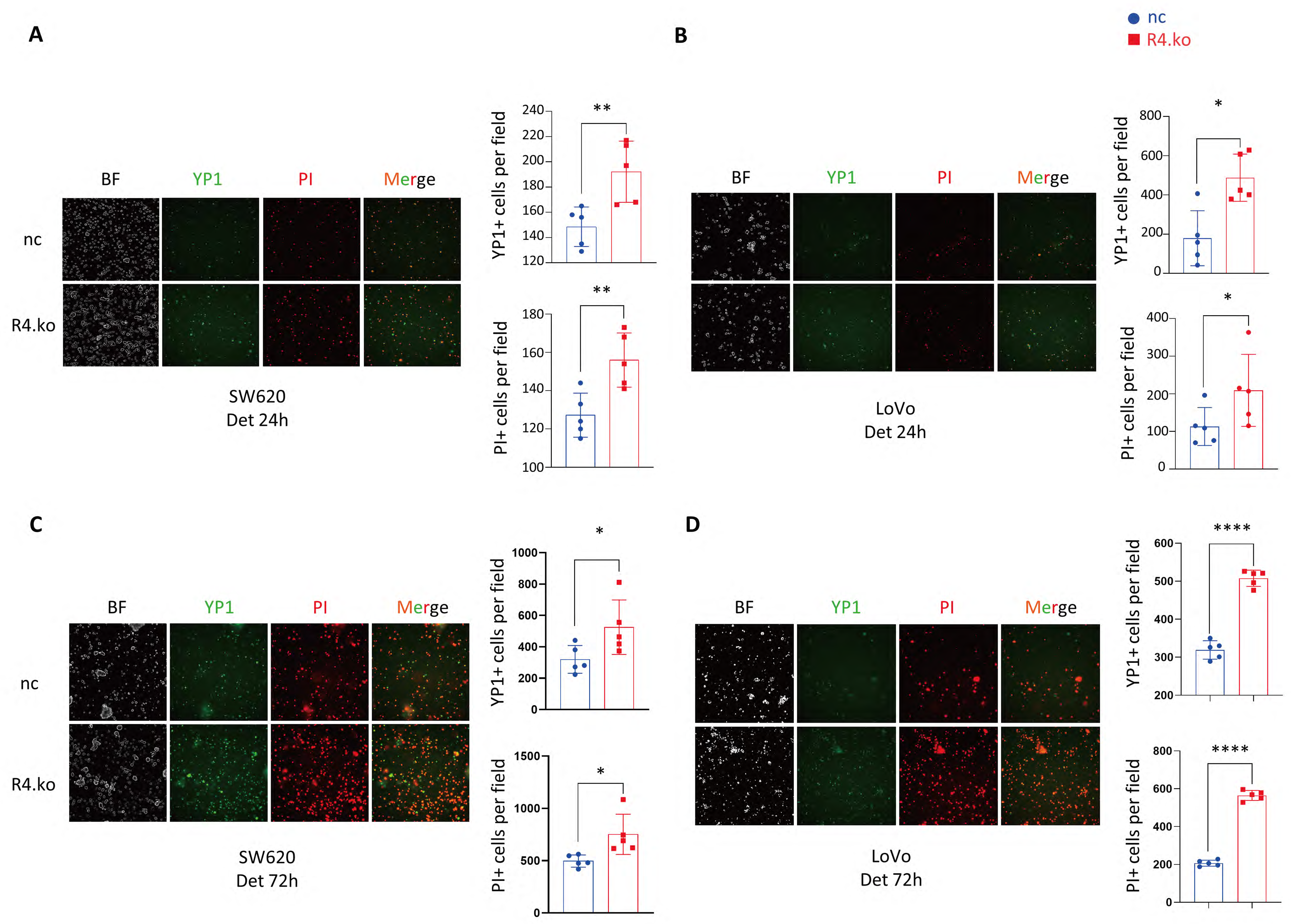
P1 and PI at 24h and 72h suspension time points. A, B. SW620 (A) and LoVo (B) after 24h suspension, in RIPK4 knockout cells, YP1 and PI signals were significantly increased, indicating that cell damage was more severe. A, B. SW620 (A) and LoVo (B) after 72h suspension. A-D. Quantitative analysis of YP1 and PI signals. The mean ± s.d., n=6, unpaired two-tailed Student‘s t-test.

**Supplementary Fig. S4.**
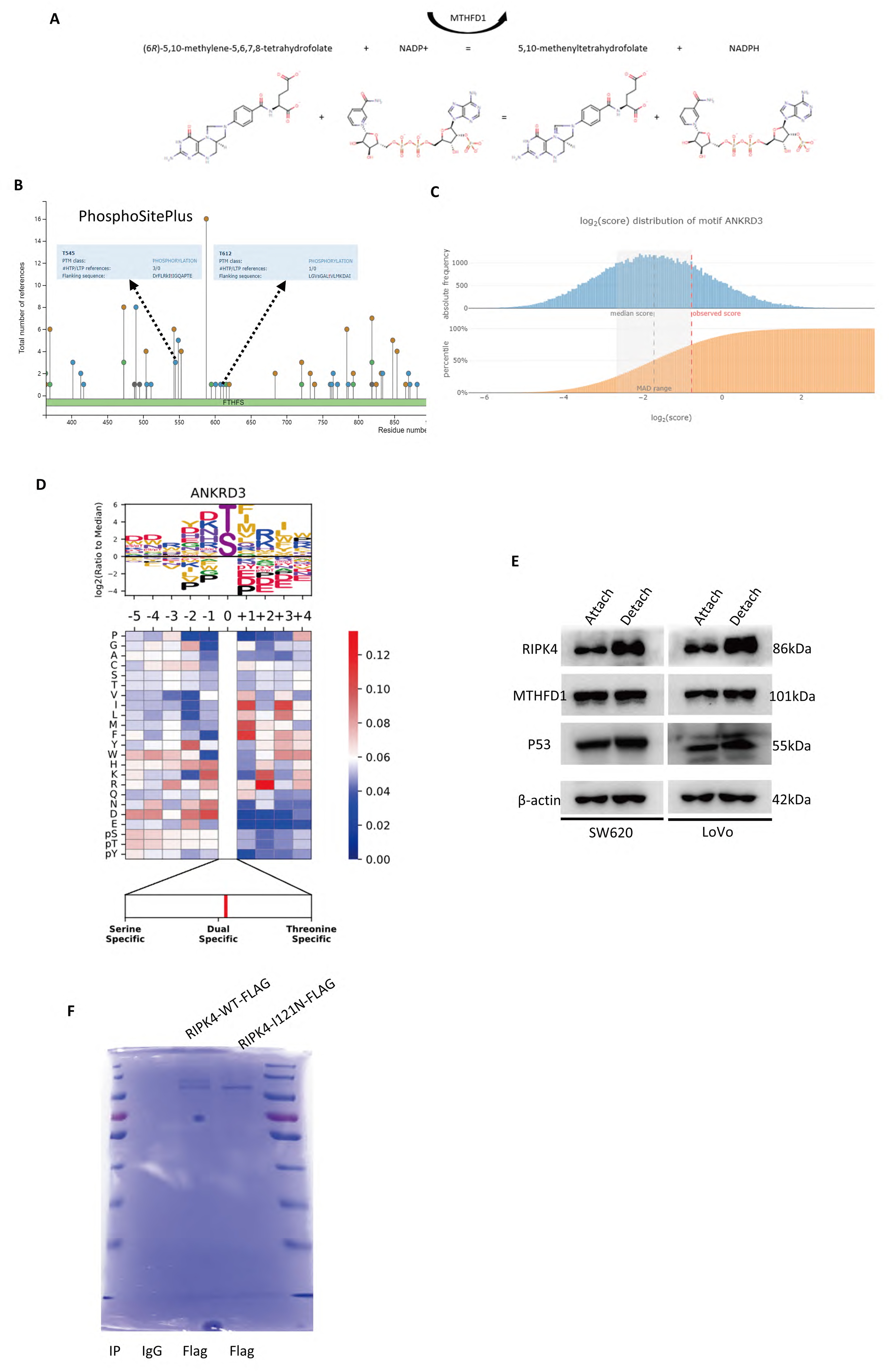
RIPK4 combines with MTHFD1 to promote threonine phosphorylation of MTHFD1. A. Functional diagram of MTHFD1 protein. MTHFD1 protein promotes the production of NADPH physiologically. B. The threonine phosphorylation at the 515-724 site of MTHFD1 protein has been previously identified. Derived from https://www.phosphosite.org. C-D. The public database predicts the effect of RIPK4 on MTHFD1 threonine phosphorylation (https://www.phosphosite.org). E. Changes in intracellular RIPK4, MTHFD1, and p53 protein levels in attach and detach states. RIPK4 and P53 protein increased during suspension, while MTHFD1 protein content did not change. F. Using Coomassie brilliant blue staining of RIPK4-WT and I121N purified from IP. The Western blot was repeated at least three times independently.

**Supplementary Fig. S5.**
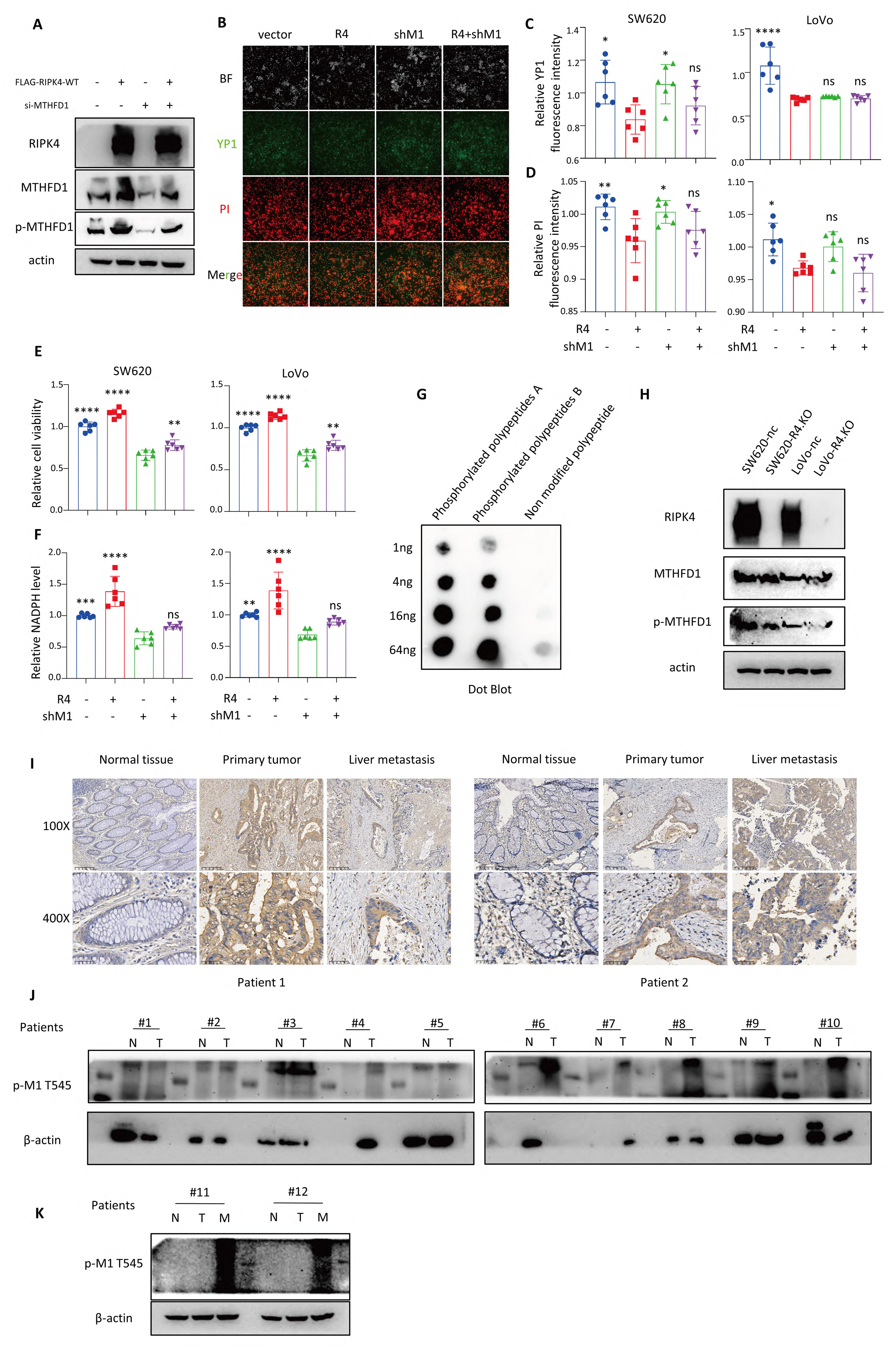
The function of RIPK4 is dependent on MTHFD1 and construction of specific phosphorylated antibody p-MTHFD1-T545. A. Changes in intracellular protein levels after overexpression of RIPK4 and/or knockdown of MTHFD1 were detected by immunoblotting. B. Changes in YP1 and PI signaling after overexpression of RIPK4 and/or knockdown of MTHFD1. C-D. Quantitative analysis of YP1 and PI signals after overexpression of RIPK4 and/or knockdown of MTHFD1. E-F. Changes in cell viability and intracellular NADPH levels after overexpression of RIPK4 and/or knockdown of MTHFD1. G. Immunospot assay of specific phosphorylated antibody p-MTHFD1-T545. H. After knocking out RIPK4, the phosphorylation changes of MTHFD1 protein were detected by p-MTHFD1-T545. I. Immunohistochemical staining of p-MTHFD1-T545 in two CRC patients with liver metastasis. J. 10 paired patient tissue protein tests p-MTHFD1-T545 K. Immunoblotting f p-MTHFD1-T545 in two CRC patients with liver metastasis. C-F. The mean ± s.d., n=6, unpaired two-tailed Student‘s t-test. A, G, H-K. The Western blot was repeated at least three times independently.

**Supplementary Fig. S6.**
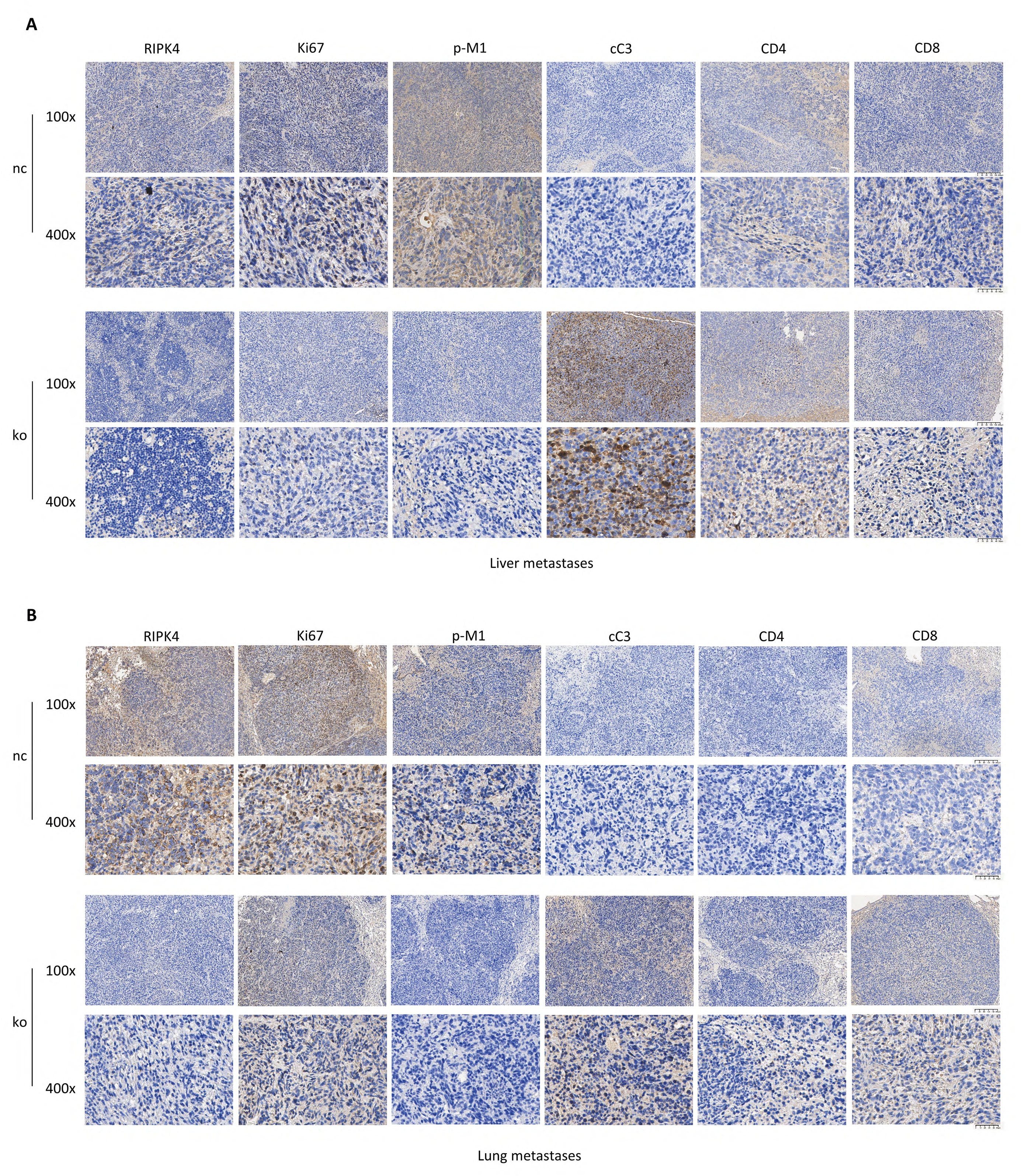
Knockout of RIPK4 reduced the viability of metastatic tumors and increased T cell infiltration. A-B. Immunohistochemical detection of liver and lung metastases in mice, including RIPK4, Ki67, p-MTHFD1-T545, cleaved caspase 3, CD4 and CD8.

## References

1. H. Brenner, M. Kloor, C. P. Pox, Colorectal cancer. Lancet 383, 1490–1502 (2014).

2. K. D. Miller, R. L. Siegel, C. C. Lin, A. B. Mariotto, J. L. Kramer, J. H. Rowland, K. D. Stein, R. Alteri, A. Jemal, Cancer treatment and survivorship statistics, 2016. CA Cancer J Clin 66, 271– 289 (2016).

3. L. Vecchione, V. Gambino, J. Raaijmakers, A. Schlicker, A. Fumagalli, M. Russo, A. Villanueva, E. Beerling, A. Bartolini, D. G. Mollevi, N. El-Murr, M. Chiron, L. Calvet, C. Nicolazzi, C. Combeau, C. Henry, I. M. Simon, S. Tian, S. in ’t Veld, G. D’ario, S. Mainardi, R. L. Beijersbergen, C. Lieftink, S. Linn, C. Rumpf-Kienzl, M. Delorenzi, L. Wessels, R. Salazar, F. Di Nicolantonio, A. Bardelli, J. van Rheenen, R. H. Medema, S. Tejpar, R. Bernards, A Vulnerability of a Subset of Colon Cancers with Potential Clinical Utility. Cell 165, 317–330 (2016).

4. S. Afrin, F. Giampieri, M. Gasparrini, T. Y. Forbes-Hernández, D. Cianciosi, P. Reboredo-Rodriguez, J. Zhang, P. P. Manna, M. Daglia, A. G. Atanasov, M. Battino, Dietary phytochemicals in colorectal cancer prevention and treatment: A focus on the molecular mechanisms involved. Biotechnol Adv 38, 107322 (2020).

5. F. Bray, J. Ferlay, I. Soerjomataram, R. L. Siegel, L. A. Torre, A. Jemal, Global cancer statistics 2018: GLOBOCAN estimates of incidence and mortality worldwide for 36 cancers in 185 countries. CA: A Cancer Journal for Clinicians 68, 394–424 (2018).

6. X. Sun, Y. Yang, X. Meng, J. Li, X. Liu, H. Liu, PANoptosis: Mechanisms, biology, and role in disease. Immunological Reviews **n/a** (2023), doi:10.1111/imr.13279.

7. J.-F. Lin, P.-S. Hu, Y.-Y. Wang, Y.-T. Tan, K. Yu, K. Liao, Q.-N. Wu, T. Li, Q. Meng, J.-Z. Lin, Z.-X. Liu, H.-Y. Pu, H.-Q. Ju, R.-H. Xu, M.-Z. Qiu, Phosphorylated NFS1 weakens oxaliplatin-based chemosensitivity of colorectal cancer by preventing PANoptosis. Sig Transduct Target Ther 7, 1–16 (2022).

8. R. Karki, B. R. Sharma, S. Tuladhar, E. P. Williams, L. Zalduondo, P. Samir, M. Zheng, B. Sundaram, B. Banoth, R. K. S. Malireddi, P. Schreiner, G. Neale, P. Vogel, R. Webby, C. B. Jonsson, T.-D. Kanneganti, Synergism of TNF-α and IFN-γ Triggers Inflammatory Cell Death, Tissue Damage, and Mortality in SARS-CoV-2 Infection and Cytokine Shock Syndromes. Cell 184, 149–168.e17 (2021).

9. A. Pandeya, T.-D. Kanneganti, Therapeutic potential of PANoptosis: innate sensors, inflammasomes, and RIPKs in PANoptosomes. Trends in Molecular Medicine 0 (2023), doi:10.1016/j.molmed.2023.10.001.

10. B. Shan, H. Pan, A. Najafov, J. Yuan, Necroptosis in development and diseases. Genes Dev. 32, 327–340 (2018).

11. D. Zhang, J. Lin, J. Han, Receptor-interacting protein (RIP) kinase family. Cell Mol Immunol 7, 243–249 (2010).

12. N. Oberbeck, V. C. Pham, J. D. Webster, R. Reja, C. S. Huang, Y. Zhang, M. Roose-Girma, S. Warming, Q. Li, A. Birnberg, W. Wong, W. Sandoval, L. G. Kőműves, K. Yu, D. L. Dugger, A. Maltzman, K. Newton, V. M. Dixit, The RIPK4-IRF6 signalling axis safeguards epidermal differentiation and barrier function. Nature 574, 249–253 (2019).

13. J. Xu, Q. Wei, Z. He, Insight Into the Function of RIPK4 in Keratinocyte Differentiation and Carcinogenesis. Front. Oncol. 10, 1562 (2020).

14. Z.-H. Qi, H.-X. Xu, S.-R. Zhang, J.-Z. Xu, S. Li, H.-L. Gao, W. Jin, W.-Q. Wang, C.-T. Wu, Q.- X. Ni, X.-J. Yu, L. Liu, RIPK4/PEBP1 axis promotes pancreatic cancer cell migration and invasion by activating RAF1/MEK/ERK signaling. Int J Oncol (2018), doi:10.3892/ijo.2018.4269.

15. J.-Y. Liu, Q.-H. Zeng, P.-G. Cao, D. Xie, X. Chen, F. Yang, L.-Y. He, Y.-B. Dai, J.-J. Li, X.-M. Liu, H.-L. Zeng, Y.-X. Zhu, L. Gong, Y. Cheng, J.-D. Zhou, J. Hu, H. Bo, Z.-Z. Xu, K. Cao, RIPK4 promotes bladder urothelial carcinoma cell aggressiveness by upregulating VEGF-A through the NF-κB pathway. Br J Cancer 118, 1617–1627 (2018).

16. S. He, X. Wang, RIP kinases as modulators of inflammation and immunity. Nat Immunol 19, 912–922 (2018).

17. E. Meylan, F. Martinon, M. Thome, M. Gschwendt, J. Tschopp, RIP4 (DIK/PKK), a novel member of the RIP kinase family, activates NF-κB and is processed during apoptosis. EMBO Rep 3, 1201–1208 (2002).

18. K. Mitchell, J. O’Sullivan, C. Missero, E. Blair, R. Richardson, B. Anderson, D. Antonini, J. C. Murray, A. L. Shanske, B. C. Schutte, R.-A. Romano, S. Sinha, S. S. Bhaskar, G. C. M. Black, J. Dixon, M. J. Dixon, Exome Sequence Identifies RIPK4 as the Bartsocas-Papas Syndrome Locus. The American Journal of Human Genetics 90, 69–75 (2012).

19. B. Poligone, E. S. Gilmore, C. V. Alexander, D. Oleksyn, K. Gillespie, J. Zhao, S. F. Ibrahim, A. P. Pentland, M. D. Brown, L. Chen, PKK Suppresses Tumor Growth and Is Decreased in Squamous Cell Carcinoma of the Skin. Journal of Investigative Dermatology 135, 869–876 (2015).

20. E. C. Cheung, K. H. Vousden, The role of ROS in tumour development and progression. Nat Rev Cancer 22, 280–297 (2022).

21. E. Madej, D. Ryszawy, A. A. Brożyna, M. Czyz, J. Czyz, A. Wolnicka-Glubisz, Deciphering the Functional Role of RIPK4 in Melanoma. Int J Mol Sci 22, 11504 (2021).

22. E. Piskounova, M. Agathocleous, M. M. Murphy, Z. Hu, S. E. Huddlestun, Z. Zhao, A. M. Leitch, T. M. Johnson, R. J. DeBerardinis, S. J. Morrison, Oxidative stress inhibits distant metastasis by human melanoma cells. Nature 527, 186–191 (2015).

23. Z. Chen, R. Tian, Z. She, J. Cai, H. Li, Role of oxidative stress in the pathogenesis of nonalcoholic fatty liver disease. Free Radic Biol Med 152, 116–141 (2020).

24. A. Chiarugi, C. Dölle, R. Felici, M. Ziegler, The NAD metabolome--a key determinant of cancer cell biology. Nat Rev Cancer 12, 741–752 (2012).

25. V. Nogueira, N. Hay, Molecular pathways: reactive oxygen species homeostasis in cancer cells and implications for cancer therapy. Clin Cancer Res 19, 4309–4314 (2013).

26. W. Xiao, R.-S. Wang, D. E. Handy, J. Loscalzo, NAD(H) and NADP(H) Redox Couples and Cellular Energy Metabolism. Antioxid Redox Signal 28, 251–272 (2018).

27. D. Xu, X. Li, F. Shao, G. Lv, H. Lv, J.-H. Lee, X. Qian, Z. Wang, Y. Xia, L. Du, Y. Zheng, H. Wang, J. Lyu, Z. Lu, The protein kinase activity of fructokinase A specifies the antioxidant responses of tumor cells by phosphorylating p62. Sci Adv 5, eaav4570 (2019).

28. W. Ying, NAD+/NADH and NADP+/NADPH in cellular functions and cell death: regulation and biological consequences. Antioxid Redox Signal 10, 179–206 (2008).

29. L. Chen, Z. Zhang, A. Hoshino, H. D. Zheng, M. Morley, Z. Arany, J. D. Rabinowitz, NADPH production by the oxidative pentose-phosphate pathway supports folate metabolism. Nat Metab 1, 404–415 (2019).

30. J. Fan, J. Ye, J. J. Kamphorst, T. Shlomi, C. B. Thompson, J. D. Rabinowitz, Quantitative flux analysis reveals folate-dependent NADPH production. Nature 510, 298–302 (2014).

31. G. S. Ducker, J. D. Rabinowitz, One-Carbon Metabolism in Health and Disease. Cell Metabolism 25, 27–42 (2017).

32. M. Yang, K. H. Vousden, Serine and one-carbon metabolism in cancer. Nat Rev Cancer 16, 650–662 (2016).

33. O. Hassin, N. B. Nataraj, M. Shreberk-Shaked, Y. Aylon, R. Yaeger, G. Fontemaggi, S. Mukherjee, M. Maddalena, A. Avioz, O. Iancu, G. Mallel, A. Gershoni, I. Grosheva, E. Feldmesser, S. Ben-Dor, O. Golani, A. Hendel, G. Blandino, D. Kelsen, Y. Yarden, M. Oren, Different hotspot p53 mutants exert distinct phenotypes and predict outcome of colorectal cancer patients. Nat Commun 13, 2800 (2022).

34. K. W. Kinzler, B. Vogelstein, Lessons from Hereditary Colorectal Cancer. Cell 87, 159–170 (1996).

35. W. A. Freed-Pastor, C. Prives, Mutant p53: one name, many proteins. Genes Dev. 26, 1268–1286 (2012).

36. M. P. Kim, G. Lozano, Mutant p53 partners in crime. Cell Death Differ 25, 161–168 (2018).

37. F. Mantovani, L. Collavin, G. Del Sal, Mutant p53 as a guardian of the cancer cell. Cell Death Differ 26, 199–212 (2019).

38. P. A. J. Muller, K. H. Vousden, Mutant p53 in Cancer: New Functions and Therapeutic Opportunities. Cancer Cell 25, 304–317 (2014).

39. M. Oren, V. Rotter, Mutant p53 Gain-of-Function in Cancer. Cold Spring Harbor Perspectives in Biology 2, a001107–a001107 (2010).

40. Q. Tang, Z. Su, W. Gu, A. K. Rustgi, Mutant p53 on the Path to Metastasis. Trends in Cancer 6, 62–73 (2020).

41. M. A. Hawk, C. L. Gorsuch, P. Fagan, C. Lee, S. E. Kim, J. C. Hamann, J. A. Mason, K. J. Weigel, M. A. Tsegaye, L. Shen, S. Shuff, J. Zuo, S. Hu, L. Jiang, S. Chapman, W. M. Leevy, R. J. DeBerardinis, M. Overholtzer, Z. T. Schafer, RIPK1-mediated induction of mitophagy compromises the viability of extracellular-matrix-detached cells. Nat Cell Biol 20, 272–284 (2018).

42. Y. Wang, Z. Zeng, J. Lu, Y. Wang, Z. Liu, M. He, Q. Zhao, Z. Wang, T. Li, Y. Lu, Q. Wu, K. Yu, F. Wang, H.-Y. Pu, B. Li, W. Jia, M. shi, D. Xie, T. Kang, P. Huang, H. Ju, R. Xu, CPT1A-mediated fatty acid oxidation promotes colorectal cancer cell metastasis by inhibiting anoikis. Oncogene 37, 6025–6040 (2018).

43. H.-L. Zhang, B.-X. Hu, Z.-L. Li, T. Du, J.-L. Shan, Z.-P. Ye, X.-D. Peng, X. Li, Y. Huang, X.-Y. Zhu, Y.-H. Chen, G.-K. Feng, D. Yang, R. Deng, X.-F. Zhu, PKCβII phosphorylates ACSL4 to amplify lipid peroxidation to induce ferroptosis. Nat Cell Biol (2022), doi:10.1038/s41556-021-00818-3.

44. H.-Q. Ju, J.-F. Lin, T. Tian, D. Xie, R.-H. Xu, NADPH homeostasis in cancer: functions, mechanisms and therapeutic implications. Sig Transduct Target Ther 5, 231 (2020).

45. W. Chaabane, S. D. User, M. El-Gazzah, R. Jaksik, E. Sajjadi, J. Rzeszowska-Wolny, M. J. Łos, Autophagy, Apoptosis, Mitoptosis and Necrosis: Interdependence Between Those Pathways and Effects on Cancer. Arch. Immunol. Ther. Exp. 61, 43–58 (2013).

46. J. Shi, W. Gao, F. Shao, Pyroptosis: Gasdermin-Mediated Programmed Necrotic Cell Death. Trends Biochem Sci 42, 245–254 (2017).

47. K. E. Christensen, C. V. Rohlicek, G. U. Andelfinger, J. Michaud, J.-L. Bigras, A. Richter, R. E. Mackenzie, R. Rozen, The MTHFD1 p.Arg653Gln variant alters enzyme function and increases risk for congenital heart defects. Hum Mutat 30, 212–220 (2009).

48. D. W. Hum, R. E. MacKenzie, Expression of active domains of a human folate-dependent trifunctional enzyme in Escherichia coli. Protein Eng 4, 493–500 (1991).

49. A. Schmidt, H. Wu, R. E. MacKenzie, V. J. Chen, J. R. Bewly, J. E. Ray, J. E. Toth, M. Cygler, Structures of three inhibitor complexes provide insight into the reaction mechanism of the human methylenetetrahydrofolate dehydrogenase/cyclohydrolase. Biochemistry 39, 6325–6335 (2000).

50. X. Huang, J. C. McGann, B. Y. Liu, R. N. Hannoush, J. R. Lill, V. Pham, K. Newton, M. Kakunda, J. Liu, C. Yu, S. G. Hymowitz, J.-A. Hongo, A. Wynshaw-Boris, P. Polakis, R. M. Harland, V. M. Dixit, Phosphorylation of Dishevelled by Protein Kinase RIPK4 Regulates Wnt Signaling. Science 339, 1441–1445 (2013).

51. A. Machado-Silva, S. Perrier, J.-C. Bourdon, p53 family members in cancer diagnosis and treatment. Seminars in Cancer Biology 20, 57–62 (2010).

52. G. Melino, p63 is a suppressor of tumorigenesis and metastasis interacting with mutant p53. Cell Death Differ 18, 1487–1499 (2011).

53. M. Muller, E. Schleithoff, W. Stremmel, G. Melino, P. Krammer, T. Schilling, One, two, three—p53, p63, p73 and chemosensitivity. Drug Resistance Updates 9, 288–306 (2006).

54. J. W. Rocco, C.-O. Leong, N. Kuperwasser, M. P. DeYoung, L. W. Ellisen, p63 mediates survival in squamous cell carcinoma by suppression of p73-dependent apoptosis. Cancer Cell 9, 45–56 (2006).

55. Y. Gong, X. Luo, J. Yang, Q. Jiang, Z. Liu, RIPK4 promoted the tumorigenicity of nasopharyngeal carcinoma cells. Biomedicine & Pharmacotherapy 108, 1–6 (2018).

56. Z. Yi, Y. Pu, R. Gou, Y. Chen, X. Ren, W. Liu, P. Dong, Silencing of RIPK4 inhibits epithelial-mesenchymal transition by inactivating the Wnt/β-catenin signaling pathway in osteosarcoma. Mol Med Report (2020), doi:10.3892/mmr.2020.10939.

57. H. Yi, Y. Su, R. Lin, X. Zheng, D. Pan, D. Lin, X. Gao, R. Zhang, Q. Guan, Ed. Downregulation of RIPK4 Expression Inhibits Epithelial-Mesenchymal Transition in Ovarian Cancer through IL-6. Journal of Immunology Research 2021, 1–15 (2021).

58. J. Li, T. McQuade, A. B. Siemer, J. Napetschnig, K. Moriwaki, Y.-S. Hsiao, E. Damko, D. Moquin, T. Walz, A. McDermott, F. K.-M. Chan, H. Wu, The RIP1/RIP3 Necrosome Forms a Functional Amyloid Signaling Complex Required for Programmed Necrosis. Cell 150, 339–350 (2012).

59. G. D. Cuny, A. Degterev, RIPK protein kinase family: Atypical lives of typical kinases. Seminars in Cell & Developmental Biology 109, 96–105 (2021).

60. T. Delanghe, Y. Dondelinger, M. J. M. Bertrand, RIPK1 Kinase-Dependent Death: A Symphony of Phosphorylation Events. Trends in Cell Biology 30, 189–200 (2020).

61. D. Xu, C. Zou, J. Yuan, Genetic Regulation of RIPK1 and Necroptosis. Annual Review of Genetics 55, 235–263 (2021).

62. N. Takahashi, H.-Y. Chen, I. S. Harris, D. G. Stover, L. M. Selfors, R. T. Bronson, T. Deraedt, K. Cichowski, A. L. Welm, Y. Mori, G. B. Mills, J. S. Brugge, Cancer Cells Co-opt the Neuronal Redox-Sensing Channel TRPA1 to Promote Oxidative-Stress Tolerance. Cancer Cell 33, 985–1003.e7 (2018).

63. K. Le Gal, M. X. Ibrahim, C. Wiel, V. I. Sayin, M. K. Akula, C. Karlsson, M. G. Dalin, L. M. Akyürek, P. Lindahl, J. Nilsson, M. O. Bergo, Antioxidants can increase melanoma metastasis in mice. Sci. Transl. Med. 7 (2015), doi:10.1126/scitranslmed.aad3740.

64. V. I. Sayin, M. X. Ibrahim, E. Larsson, J. A. Nilsson, P. Lindahl, M. O. Bergo, Antioxidants Accelerate Lung Cancer Progression in Mice. Sci. Transl. Med. 6 (2014), doi:10.1126/scitranslmed.3007653.

65. J. M. Ubellacker, A. Tasdogan, V. Ramesh, B. Shen, E. C. Mitchell, M. S. Martin-Sandoval, Z. Gu, M. L. McCormick, A. B. Durham, D. R. Spitz, Z. Zhao, T. P. Mathews, S. J. Morrison, Lymph protects metastasizing melanoma cells from ferroptosis. Nature 585, 113–118 (2020).

66. P. Lee, S. Jiang, Y. Li, J. Yue, X. Gou, S. Chen, Y. Zhao, M. Schober, M. Tan, X. Wu, Phosphorylation of Pkp1 by RIPK 4 regulates epidermal differentiation and skin tumorigenesis. EMBO J 36, 1963–1980 (2017).

67. H. Yu, H. Wang, H.-R. Xu, Y.-C. Zhang, X.-B. Yu, M.-C. Wu, G.-Z. Jin, W.-M. Cong, Overexpression of MTHFD1 in hepatocellular carcinoma predicts poorer survival and recurrence. Future Oncol 15, 1771–1780 (2019).

68. S. Agarwal, M. Behring, K. Hale, S. Al Diffalha, K. Wang, U. Manne, S. Varambally, MTHFD1L, A Folate Cycle Enzyme, Is Involved in Progression of Colorectal Cancer. Transl Oncol 12, 1461–1467 (2019).

69. K. Ding, J. Jiang, L. Chen, X. Xu, Methylenetetrahydrofolate Dehydrogenase 1 Silencing Expedites the Apoptosis of Non-Small Cell Lung Cancer Cells via Modulating DNA Methylation. Med Sci Monit 24, 7499–7507 (2018).

70. D. Lee, I. M.-J. Xu, D. K.-C. Chiu, R. K.-H. Lai, A. P.-W. Tse, L. Lan Li, C.-T. Law, F. H.-C. Tsang, L. L. Wei, C. Y.-K. Chan, C.-M. Wong, I. O.-L. Ng, C. C.-L. Wong, Folate cycle enzyme MTHFD1L confers metabolic advantages in hepatocellular carcinoma. Journal of Clinical Investigation 127, 1856–1872 (2017).

71. Q. Meng, Y.-X. Lu, C. Wei, Z.-X. Wang, J.-F. Lin, K. Liao, X.-J. Luo, K. Yu, Y. Han, J.-J. Li, Y.-T. Tan, H. Li, Z.-L. Zeng, B. Li, R.-H. Xu, H.-Q. Ju, Arginine methylation of MTHFD1 by PRMT5 enhances anoikis resistance and cancer metastasis. Oncogene 41, 3912–3924 (2022).

72. W. Sang, Y. Zhou, H. Chen, C. Yu, L. Dai, Z. Liu, L. Chen, Y. Fang, P. Ma, X. Wu, H. Kong, W. Liao, H. Jiang, J. Qian, D. Wang, Y.-H. Liu, Receptor-interacting protein kinase 2 is an immunotherapy target in pancreatic cancer. Cancer Discovery (2023), doi:10.1158/2159-8290.CD-23-0584.

73. B. S. Tan, K. H. Tiong, H. L. Choo, F. F.-L. Chung, L.-W. Hii, S. H. Tan, I. K. S. Yap, S. Pani, N. T. W. Khor, S. F. Wong, R. Rosli, S.-K. Cheong, C.-O. Leong, Mutant p53-R273H mediates cancer cell survival and anoikis resistance through AKT-dependent suppression of BCL2-modifying factor (BMF). Cell Death Dis 6, e1826 (2015).

74. G. Xiao, D. Lundine, G. K. Annor, J. Canar, V. Ellison, A. Polotskaia, P. L. Donabedian, T. Reiner, G. F. Khramtsova, O. I. Olopade, A. Mazo, J. Bargonetti, Gain-of-Function Mutant p53 R273H Interacts with Replicating DNA and PARP1 in Breast Cancer. Cancer Res 80, 394–405 (2020).

